# Bma-LAD-2, an intestinal cell adhesion protein, as a potential therapeutic target for lymphatic filariasis

**DOI:** 10.1101/2021.09.09.459564

**Authors:** Alexander F. Flynn, Rebekah T. Taylor, Marzena E. Pazgier, Sasisekhar Bennuru, Alyssa R. Lindrose, Spencer L. Sterling, C. Paul Morris, Tim K. Maugel, Thomas B. Nutman, Edward Mitre

## Abstract

Lymphatic filariasis (LF) is a debilitating disease that afflicts over 70 million people worldwide. It is caused by the parasitic nematodes *Wuchereria bancrofti*, *Brugia malayi*, and *Brugia timori*. While efforts to eliminate LF have seen substantial success, complete eradication will likely require more time and resources than predicted. Identifying new drug and vaccine targets in adult filariae could help elimination efforts.

This study’s aim was to evaluate intestinal proteins in adult *Brugia malayi* worms as possible therapeutic targets. Using siRNA, we successfully inhibited transcripts of four candidate genes: Bma-Serpin, Bma-ShTK, Bma-Reprolysin, and Bma-LAD-2. Of those, Bma-LAD-2, an immunoglobulin superfamily cell adhesion molecule (IgSF CAM), was determined to be essential for adult worm survival. We observed a 70.42% knockdown in Bma-LAD-2 transcript levels 1 day post-siRNA incubation and an 87.02% reduction in protein expression 2 days post-siRNA incubation. This inhibition of Bma-LAD-2 expression resulted in an 80% decrease in worm motility over 6 days, a 93.43% reduction in microfilaria release (Mf) by day 6 post-siRNA incubation, and a significant decrease in MTT reduction. Transmission electron microscopy revealed the loss of microvilli and unraveling of mitochondrial cristae in the intestinal epithelium of Bma-LAD-2 siRNA-treated worms. Strikingly, Bma-LAD-2 siRNA-treated worms exhibited an almost complete loss of pseudocoelomic fluid, suggesting that loss of these tight junctions led to the leakage and subsequent loss of the worm’s structural integrity. Luciferase immunoprecipitation system assay demonstrated that serum from 30 patients with LF did not have detectable IgE antibodies against Bma-LAD-2, indicating that LF exposure does not result in IgE sensitization to this antigen.

These results indicate that Bma-LAD-2 is an essential protein for adult *Brugia malayi* and may be an effective drug or vaccine target. In addition, these findings further validate the strategy of targeting the worm intestine to prevent and treat helminthic infections.

**Author Summary:** *Brugia malayi* is a parasitic nematode that can cause lymphatic filariasis, a debilitating disease prevalent in tropical and subtropical countries. Significant progress has been made towards eliminating the disease. However, complete eradication may require new therapeutics such as drugs or a vaccine that kill adult filariae. In this study, we identified an immunoglobulin superfamily cell adhesion molecule (Bma-LAD-2) as a potential drug and vaccine candidate. When we knocked down Bma-LAD-2 expression, we observed a decrease in worm motility, fecundity, and metabolism. We also visualized the loss of microvilli, destruction of the mitochondria in the intestinal epithelium, and loss of pseudocoelomic fluid contents after Bma-LAD-2 siRNA treatment. Finally, we demonstrated that serum from filaria-infected patients does not contain preexisting IgE to Bma-LAD-2, which indicates that this antigen would likely be safe to administer as a vaccine in endemic populations.

## Introduction

Over 70 million people are infected worldwide with lymphatic filariasis (LF), a debilitating disease characterized by severe lymphedema, elephantiasis, and hydrocele [1, 2]. LF is caused by the parasitic nematodes *Wuchereria bancrofti, Brugia malayi,* and *Brugia timori*. Currently, efforts to eliminate this disease have been spearheaded by the Global Programme to Eliminate Lymphatic Filariasis (GPELF) [3]. While this campaign has reduced the overall prevalence of the disease, elimination target dates have been difficult to meet. According to a January 2020 WHO report on ending neglected tropical diseases, of the 71 countries endemic for LF in 2000, only 17 have been declared free of LF as a public health problem. The original goal set by the GPELF called for global elimination of LF as a public health problem by 2020, but this WHO report established a new goal of eliminating LF as a public health problem from 81% of endemic countries by 2030 [4]. New strategies and therapeutics would likely improve our ability to meet this new target [5–7].

Current therapies for LF include diethylcarbamazine (DEC), ivermectin (IVM), and albendazole. While triple drug therapy with all three of these agents has shown great promise [8, 9], a major limitation of these medications is that DEC and IVM cannot be administered empirically in areas endemic for *Loa loa* or *Onchocerca volvulus* because the drugs can precipitate severe side effects by rapid killing of Mf [10–13].

To avoid side effects from killing of microfilariae in co-endemic populations and to potentially enable a single treatment cure of filarial infections, our group has focused on identifying drug and/or vaccine targets specific to adult filarial worms. Because adult worms contain a complete intestinal tract, whereas microfilariae do not, our group evaluated the intestinal tract of adult filarial worms as a possible source of therapeutic targets. Already, this strategy appears to be promising against other helminths. Numerous studies have demonstrated protection against hookworm and barber pole worm infection using nematode intestinal antigens as vaccine candidates [14–19]. Furthermore, there seems to be little specific IgE against intestinal antigens in the sera of infected animal models as well as in previously exposed individuals [20, 21], suggesting that intestinal antigens maybe safe to administer as vaccines in endemic areas.

Our lab previously performed a proteomic analysis of the body wall, gut, and reproductive tract of *Brugia* adult worms [22]. We identified 396 proteins specific for the intestine, and then selected 9 for evaluation as potential drug and therapeutic targets. The selection criteria were 1) having high homology with orthologs in other filarial species and low homology to humans, 2) a large extracellular domain potentially accessible to drugs and antibody, and, 3) a predicted function likely essential for adult filaria survival. Previous work we have conducted found that another filarial intestinal antigen, Bm-UGT (UDP-glucuronosyl transferase), was essential for adult *B. malayi* survival and could be targeted with probenecid to achieve death of adult worms [23].

Using siRNA inhibition, we successfully knocked-down 4 target proteins. Of these, Bma-LAD-2, an IgSF CAM, was found to be essential for adult worm survival. Suppression of Bma-LAD-2 expression resulted in decreased worm motility, metabolism, and Mf release. Electron microscopy revealed that inhibition of Bma-LAD-2 resulted in almost complete loss of pseudocoelomic fluid, suggesting that disrupting the tight junctions between filarial intestinal cells and causing subsequent disruption of the worms’ “hydrostatic skeleton” may be a novel mechanism to kill filarial parasites.

## Results

### Structural Analysis of Bma-LAD-2

The Bma-LAD-2 protein is 1171 AA in length (MW of 133310.4 Da), with a signal peptide, AA 1-18, a large extracellular segment at position 19-1120, a transmembrane portion at AA 1121-1143, and a small cytoplasmic domain at position 1143-1171 (Fig S1). The putative domain organization and model of the structure of the extracellular domain (residues 18-1120) is shown in Fig 1 for both the Bma-LAD-2 monomer and dimer. The Bma-LAD-2 monomer is predicted to fold into 6 immunoglobulin domains (Ig1-Ig6) followed by 5 fibronectin-type domains (FN1-5) (Fig 1A). The outermost N-terminal Ig domains are predicted to homodimerize to form tight junctions. The Bma-LAD-2 dimer model, based on dimerization mode of the homologous protein neurofascin [24], is stabilized by contacts between the domains of Ig1 and Ig2 paired in an orthogonal side-to-side stacking mode (Fig 1B). It is likely that prevention and/or disruption of formation of the tight Ig junction or destabilization and/or disruption of the Bma-LAD-2 dimer may lead to loss of Bma-LAD-2 function.

**Fig 1.**
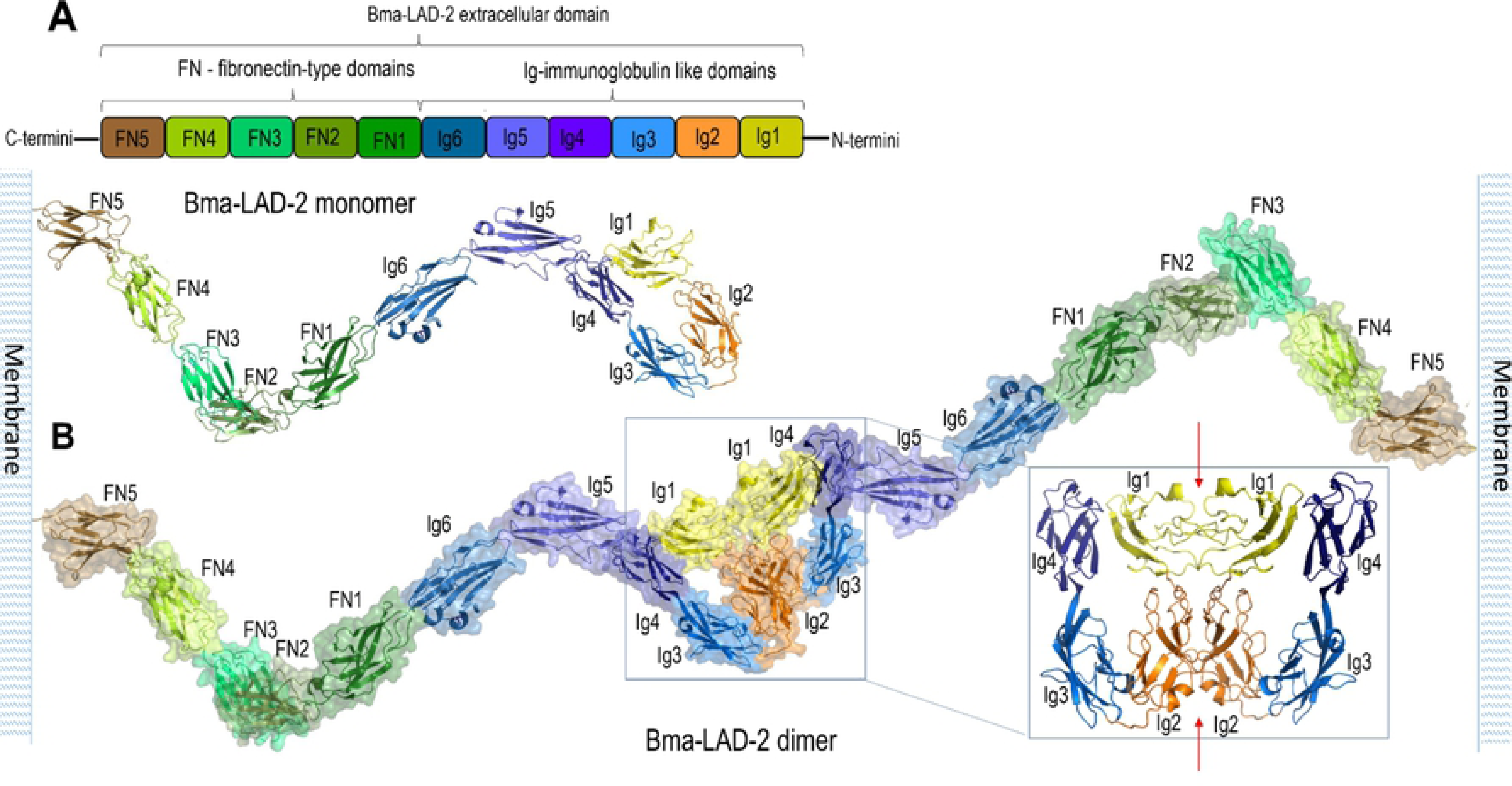
**Molecular organization of Bma-LAD-2 extracellular domain**. (A) Bma-LAD-2 monomer. Schematic domain organization (top panel) and model of monomer structure (bottom panel) assembled based on sequence similarity and available crystal structures of homologous proteins as described in Material and Methods. (B) Putative structure of Bma-LAD-2 dimer. Expanded view shows the dimer interface (indicated by red arrows) with Ig domains as labeled.

### Bma-LAD-2 is phylogenetically related to orthologs found in other filarial worms

Bma-LAD-2 has previously been shown to be a protein localized to the gut of adult *B. malayi* worms and to have a high predicted sequence homology with other filarial orthologs [22]. In this study, we generated a phylogenetic tree (Fig 2) to view the level of evolutionary relatedness between Bma-LAD-2 and orthologs from other filarial species and helminths. We found a close phylogenetic relation between Bma-LAD-2 and orthologs found in other filarial species and with orthologs of the intestinal helminths. Furthermore, the large phylogenetic distance to orthologs in humans, dogs, and cats suggests that filarial protein can likely be targeted by medications or vaccines with little risk to the host.

**Fig 2.**
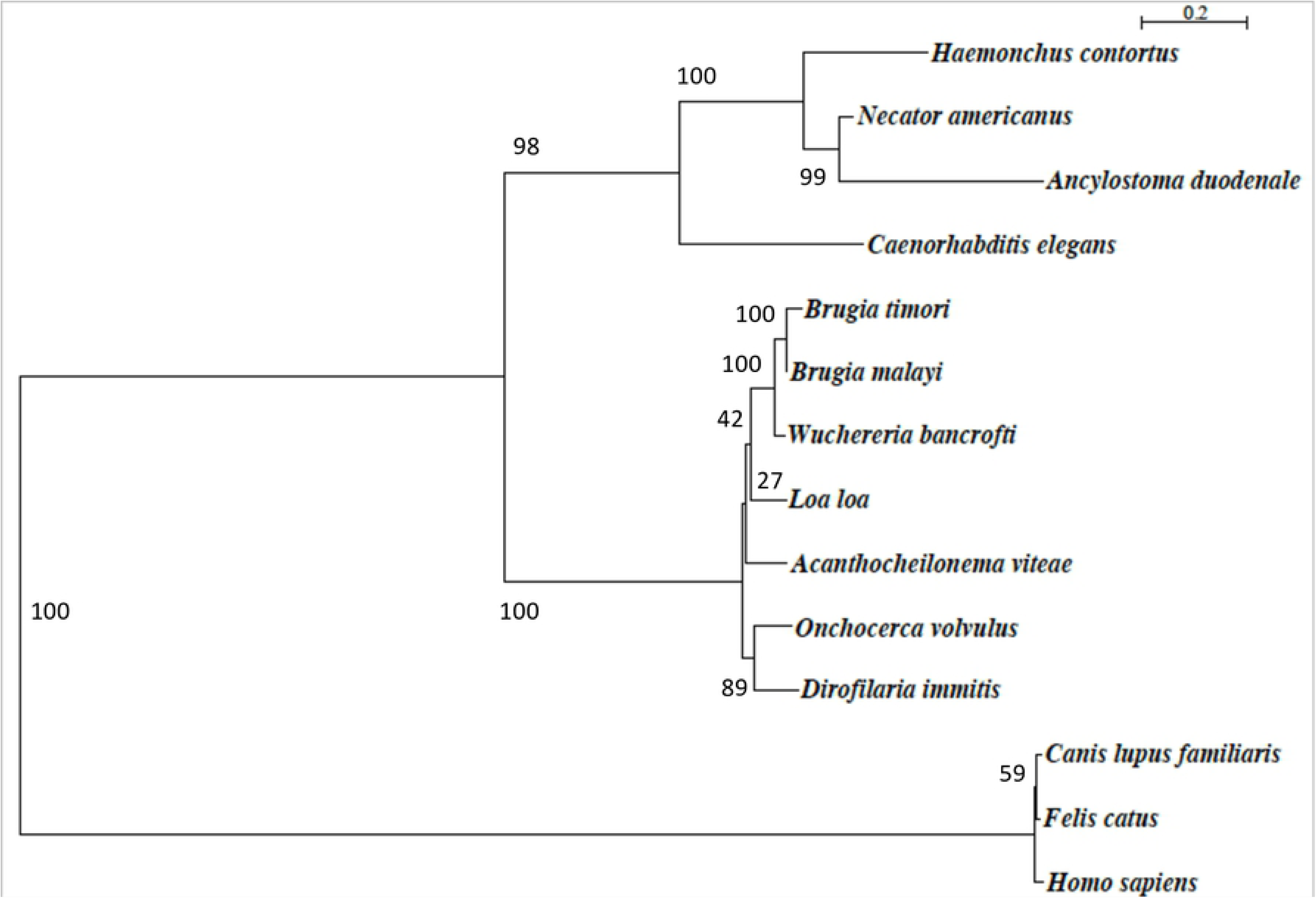
**Phylogenetic tree of *Bma-*LAD-2 and orthologs from other helminths.** The amino acid sequence of *Bma*-LAD-2 (WormBase gene ID: WBGene00227085) has high level of relatedness to other filarial species and is evolutionarily distant from cats, dogs, and humans. Based on sequence alignment using MUltiple Sequence Comparison by Log-Expectation (MUSCLE), the phylogenetic tree was constructed by the maximum likelihood method. The phylogenetic scale represents genetic change as defined by the average number of nucleotide substitutions per site. The numbers at each branch represent the bootstrap value out of a 100.

### Bma-LAD-2 is expressed throughout the lifecycle of *B. malayi* adult worms

To determine whether Bma-LAD-2 is expressed in a stage-specific manner, we analyzed stage-specific transcriptomic data on *Brugia malayi* worms [25]. We found that Bma-LAD-2 RNA was expressed in the Mf, L3, L4, and adult stages regardless of gender. The highest expression levels based on normalized read values occured in mature microfilaria (44 RPKM) followed by adult female filaria (26 RPKM). Overall, Bma-LAD-2 transcript levels appeared to be similar across the different life stages. Next, we looked at a study evaluating the proteome for the different stages of *B. malayi* [26]. The study matched 3 unique peptides to Bma-LAD-2 from Mf, 1 from the L3 stage, 2 from adult females, and 1 from adult males, suggesting that Bma-LAD-2 is expressed during Mf and L3 stages, as well as in both genders of adult worms. Like the transcriptome data, these spectra values indicate fairly consistent expression of Bma-LAD-2 among the larval stages.

### Cy3-labeled siRNA enters the intestinal tract of *B. malayi* adult worms

Prior to siRNA knockdown, we investigated whether Bma-LAD-2 siRNA conjugated to cy3 could be visualized in the intestinal tract of adult filariae. We incubated the adult female worms with labeled siRNA for 24 hrs and then observed them using epifluorescent microscopy. The cy3-labeled siRNA (Fig 3A-B) was easily seen in the intestinal tract of the treated worms. As expected, minimal signal was visualized in the intestine of worms treated with only culture media (Fig 3C-D). We therefore concluded that siRNA targeting Bma-LAD-2 transcript could enter the intestinal tract of adult filarial worms.

**Fig 3.**
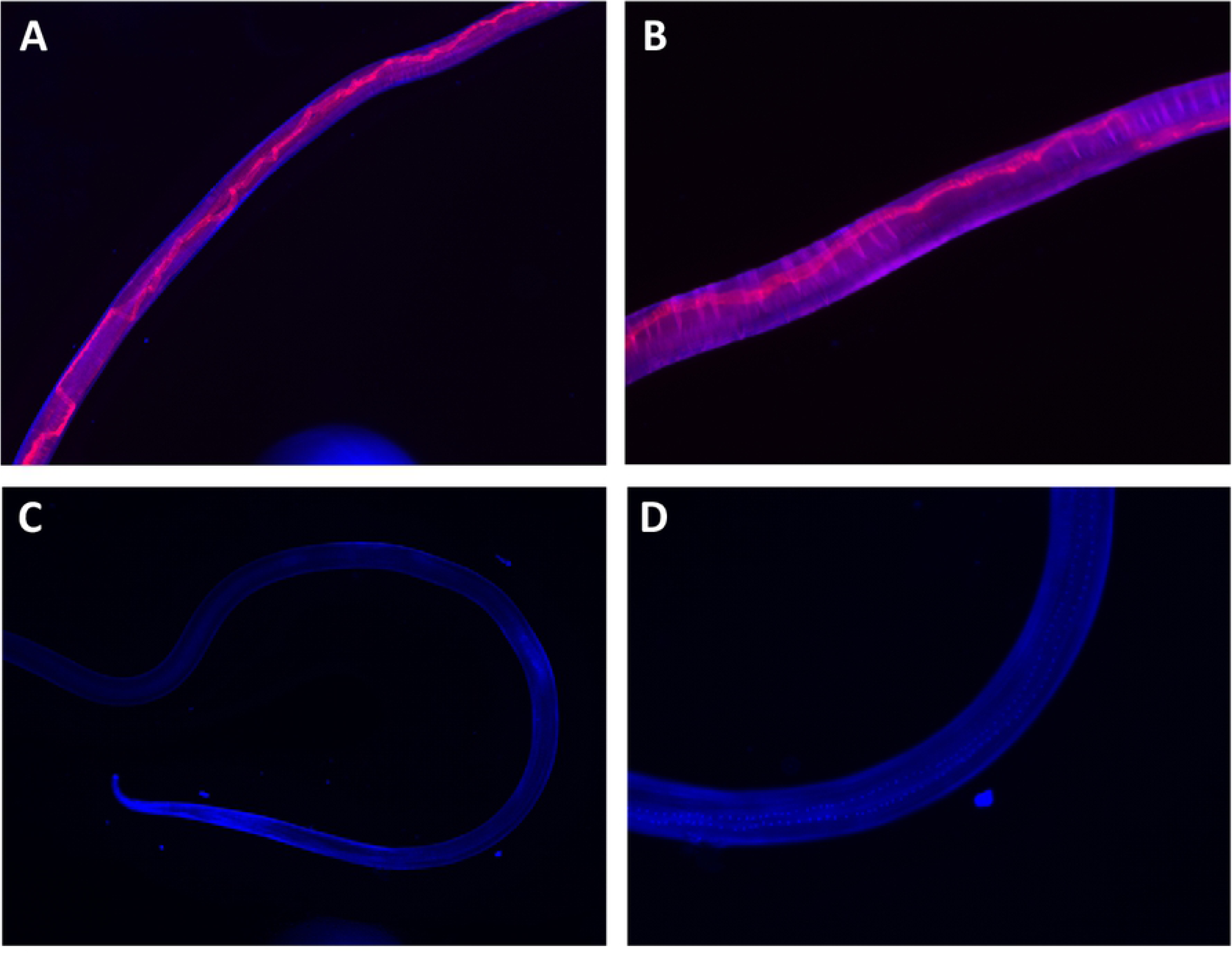
**cy3-labeled *Bma*-LAD-2 siRNA entry into the intestinal tract of female *B. malayi* adult worms.** *B. malayi* adult female worms soaked in cy3-labeled siRNA (red) for 24 hrs and visualized at magnifications of (A) 40x and (B) 100x. As a negative control, worms were cultured in only media for 24 hrs and visualized at magnifications of (C) 40x and (D) 100x. Worms were counterstained with DAPI (blue).

### Bma-LAD-2 siRNA treatment results in reduced transcript and protein expression

After observation of Bma-LAD-2 siRNA entry into the intestine, gene and protein expression was evaluated by quantitative reverse transcription PCR (RT-qPCR) and Western blot respectively. cDNA was generated using mRNA isolated from adult female *B. malayi* cultured in media alone, Bma-LAD-2 siRNA, and scrambled siRNA for 1 day and 6 days post-siRNA incubation. By quantifying *B. malayi lad-2* gene expression normalized to *Bma-gapdh* under each condition, *we* observed a 70.42% decrease in Bma-LAD-2 transcript levels (Fig 4A) in worms treated with target specific siRNA (mean=0.2662) compared to the scrambled siRNA-treated filariae (mean=0.9) relative to the media control group at 1 day post-siRNA incubation. Bma-LAD-2 protein expression was visualized by immunoblotting in worms treated with specific or scrambled siRNA. A dramatic decrease in Bma-LAD-2 expression was observed in the specific siRNA-treated worms compared to the controls (Fig 4B).

**Fig 4.**
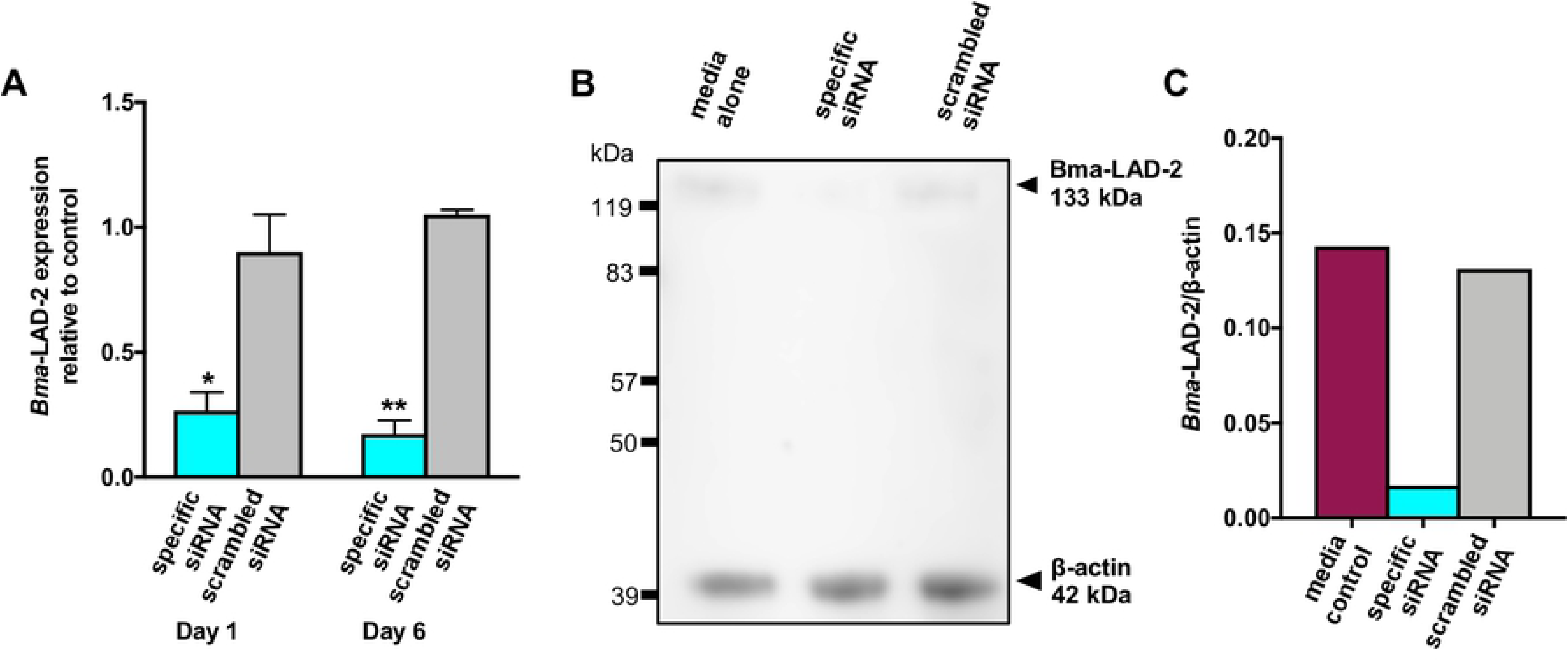
**Treatment with Bma*-*LAD-2*-*specific siRNA reduces target transcript and protein levels in adult female *B. malayi* worms.** Adult female filaria were cultured in Bma*-*LAD-2 siRNA, scrambled siRNA, or media alone. (A) The *Bma-lad-2* transcript level was reduced in specific siRNA-treated groups (n=3) compared to the scrambled controls (n=3) normalized to *Bma-gapdh*. Ordinary one-way ANOVA followed by Tukey’s multiple comparison test was used to generate the p-values. These experiments were successfully repeated twice and the data shown is from a single representative experiment; mean + SEM. (B) Bma-LAD-2 levels were assessed 24 hrs post-siRNA incubation by Western blot using anti-Bma-LAD-2 antibodies. (C) Western blot quantification was performed using the ImageStudioLite software. The signal intensities of anti-Bma-LAD-2 were normalized to those of beta-actin.

### Reduced worm viability and fecundity in Bma-LAD-2 siRNA-treated adult filariae

We evaluated the effects of Bma-LAD-2 knockdown on worm motility, Mf release, and (4,5-dimethylthiazol-2-yl)-2,5-diphenyltetrazolium bromide (MTT) reduction. At day 1 post-siRNA incubation, worm motility (Fig 5A) was significantly reduced (p<0.0001) in worms soaked in Bma-LAD-2 siRNA (mean=0.8) compared to the control group (mean=4). By day 6 post-incubation, we observed an 85% reduction in motility relative to the controls (mean=4) for the specific siRNA-treated group (mean=0.6).

**Fig 5.**
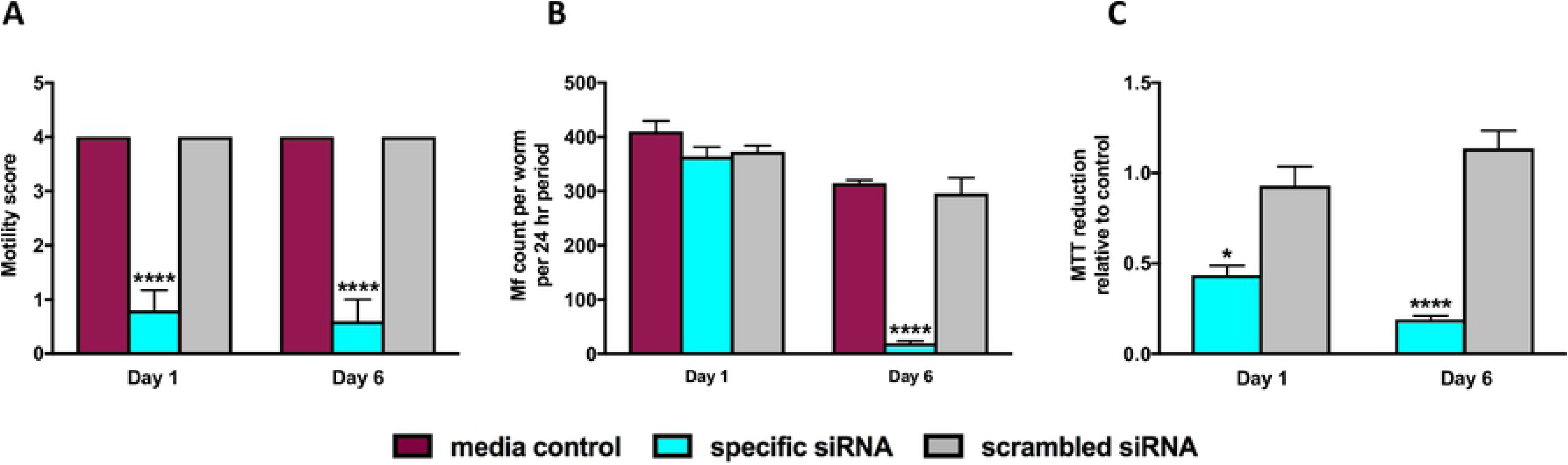
**Bma*-*LAD-2 knockdown results in decreased worm motility, microfilariae release, and metabolism.** Reducing Bma*-*LAD-2 expression in female *B. malayi* adult worms resulted in decreased (A) motility (n=5; Day 1, p<0.0001; Day 6, p=0.0004), (B) microfilaria count per worm (n=5; Day 1, p=ns; Day 6, p<0.0001) per 24-hr period, and (C) metabolism (n=2; Day 1, p=0.0139; Day 6, p=0.0007) as measured by MTT reduction at timepoints 1 and 6 days post-siRNA treatment. For motility (A) and microfilaria release (B), an ordinary one-way ANOVA followed by Tukey’s multiple comparison test was used to generate the p-values while p-values for metabolism (C) were generated by an unpaired t-test. These experiments were successfully repeated twice and the data presented is representative of a single experiment; mean + SEM.

We next evaluated Mf release per adult worm per 24 hr period for each group at timepoints 1 and 6 days post-siRNA incubation. While no reduction in Mf release was observed at one day after treatment with siRNA, Mf release was 93.4% lower from Bma-LAD-2 siRNA treated adult filariae compared to Mf release from worms incubated with media alone (Fig 5B).

Finally, two randomly selected adult worms from each group were assessed by MTT reduction assay at each timepoint. At day 1 post-siRNA treatment, MTT reduction by *B. malayi* treated with target siRNA was 46% less than MTT reduction observed by worms treated with scrambled siRNA relative to the media control group (p=0.014, Fig 5C). By day 6, we observed an 83.25% decrease in MTT reduction by the Bma-LAD-2 siRNA group compared to worms treated with scrambled siRNA (Fig 5C). Given the above results, we conclude that Bma-LAD-2 is an essential protein for *B. malayi* adult worm survival *in vitro* and knockdown results in death of the adult worm as defined by motility, fecundity, and metabolism.

### Bma-LAD-2 knockdown results in ablation of microvilli and loss of pseudocoelomic fluid

Bma-LAD-2 is predicted to be an adhesion protein located at the apical junction of the intestinal tract [22]. Therefore, we evaluated the structure of the adult filaria intestinal tract after treatment with Bma-LAD-2 siRNA using transmission electron microscopy (TEM). As seen in Fig 6, microvilli lining the epithelium of the intestinal tract are present in untreated *B. malayi* adults. In contrast, adult worms treated with Bma-LAD-2 siRNA exhibit near complete ablation of intestinal microvilli (Fig 7). In addition, many of the apical junctions in the intestinal tract of treated worms appear shortened, and many of the mitochondria had misshaped cristae.

**Fig 6.**
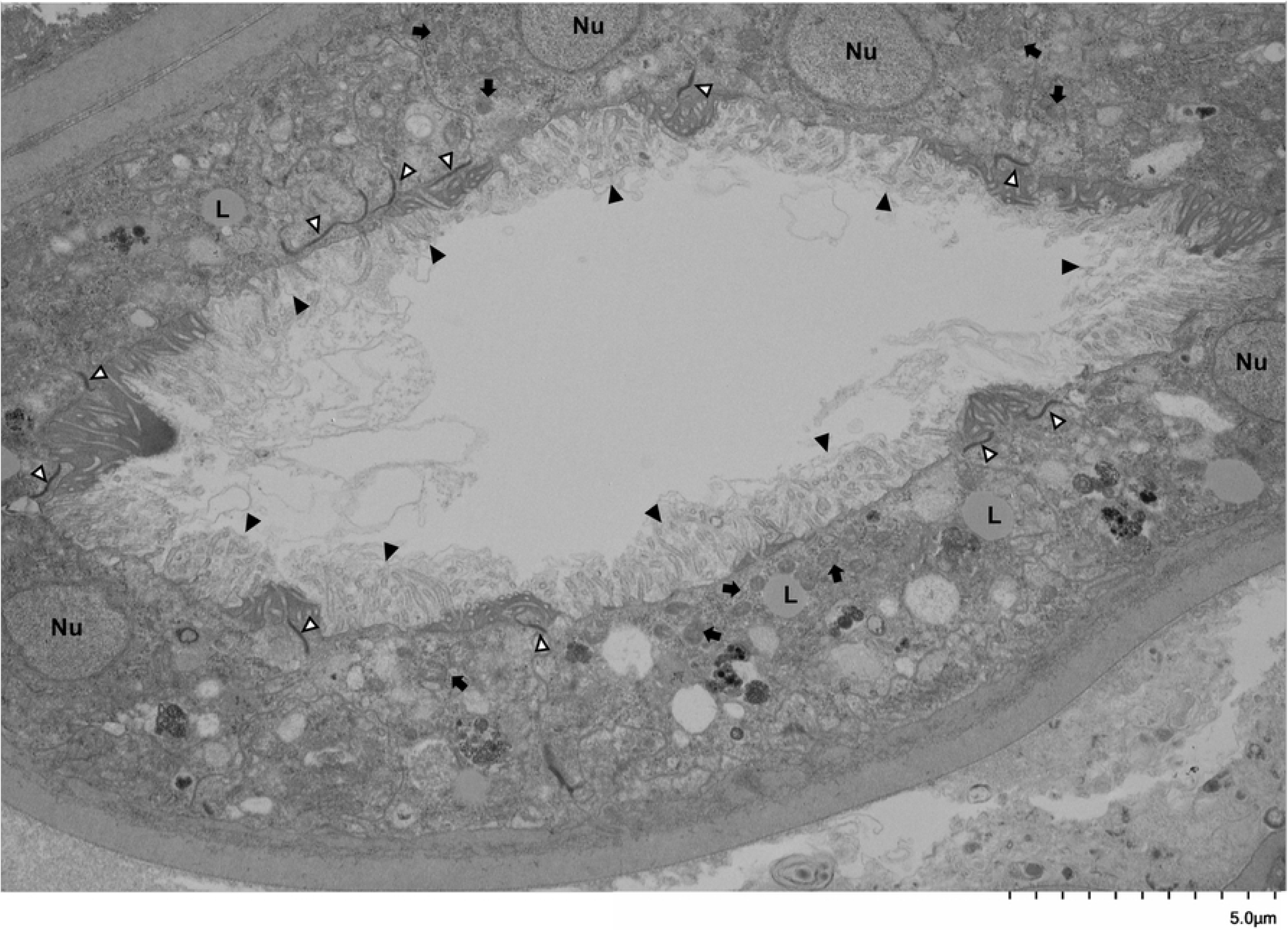
**Intestinal tract of a female *B. malayi* adult worm.** Image was captured by transmission electron microscopy (TEM) at 4,000x. Adult filaria were cultured in media alone for 72 hrs. Microvilli (black arrowhead) line the apical surface of the intestinal epithelium. Other structures visible are apical junctions (white arrowhead), nuclei (Nu), lipid droplets (L), and mitochondria (black arrow)

**Fig 7.**
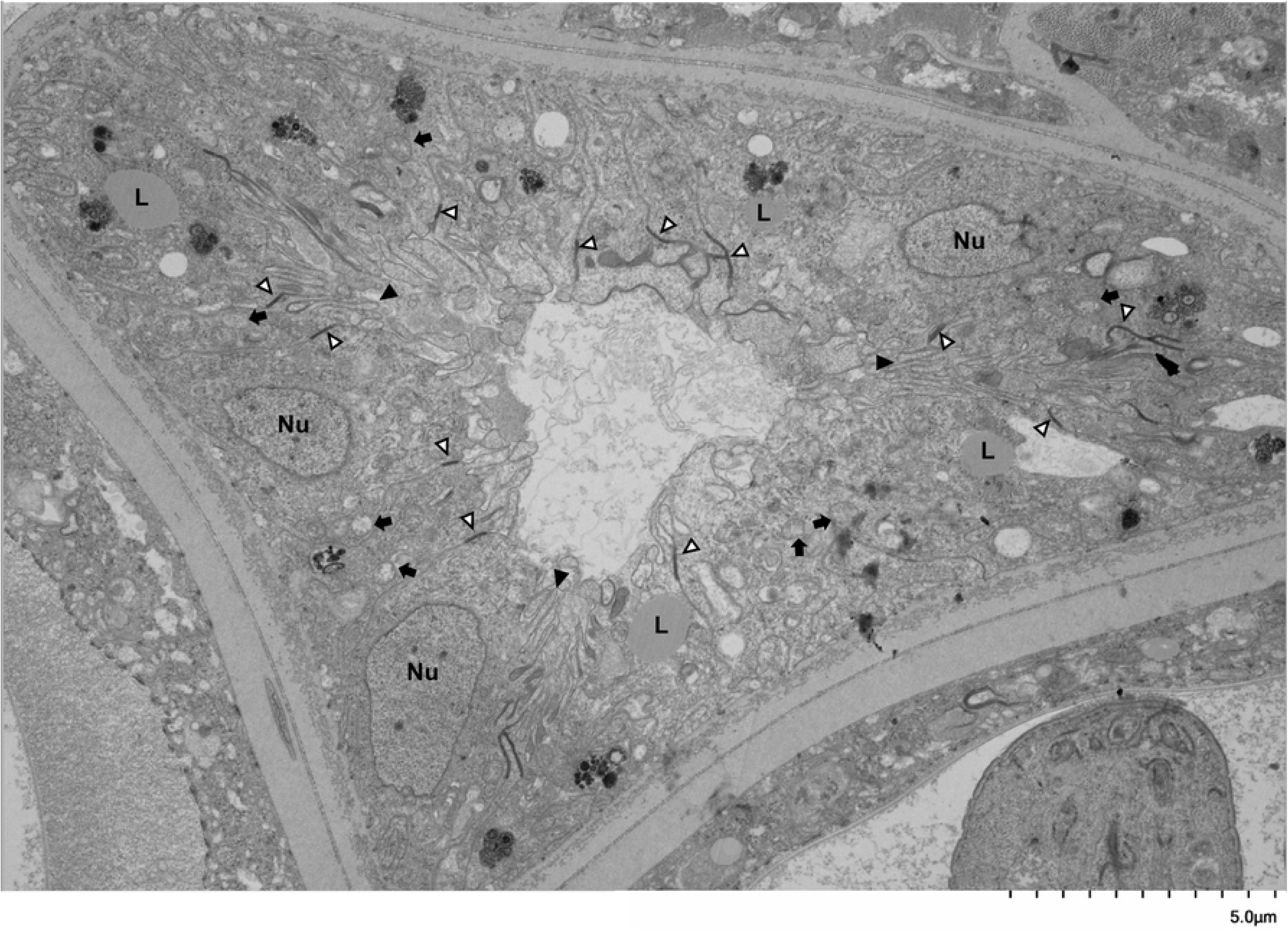
**Intestinal tract of a female *B. malayi* adult worm treated with *Bma-*LAD-2 siRNA.** Image was captured by transmission electron microscopy (TEM) at 4,000x. Adult filaria were incubated with *Bma-*LAD-2 siRNA for 24 hrs and then cultured in media alone for an additional 48 hrs. Microvilli (black arrowhead) are largely absent from the apical surface of the intestinal epithelium. There are some vestigial microvilli have been invaginated by the surrounding epithelium. Other structures visible are apical junctions (white arrowhead), nuclei (Nu), and lipid droplets (L). Many mitochondria (black arrow) appear to have unraveled cristae.

Interestingly, when observed at lower magnification, enabling visualization of the entire nematode cross-section, it is apparent that the pseudocoelomic fluid is absent or markedly diminished in treated adult filariae (Fig 8). While untreated worms display pseudocoelomic fluid separating the intestinal tract from the uterine tubes at all cross-sections analyzed, Bma-LAD-2 siRNA-treated worms demonstrated direct contact between the intestinal tract and the uterine tubes. In toto, the imaging findings suggest that knockdown of Bma-LAD-2 results in reduction of tight junctions between intestinal epithelial cells, escape of pseudocoelomic fluid from the internal body cavity into the intestinal tract, and subsequent loss of the integrity of the filarial hydroskeleton.

**Fig 8.**
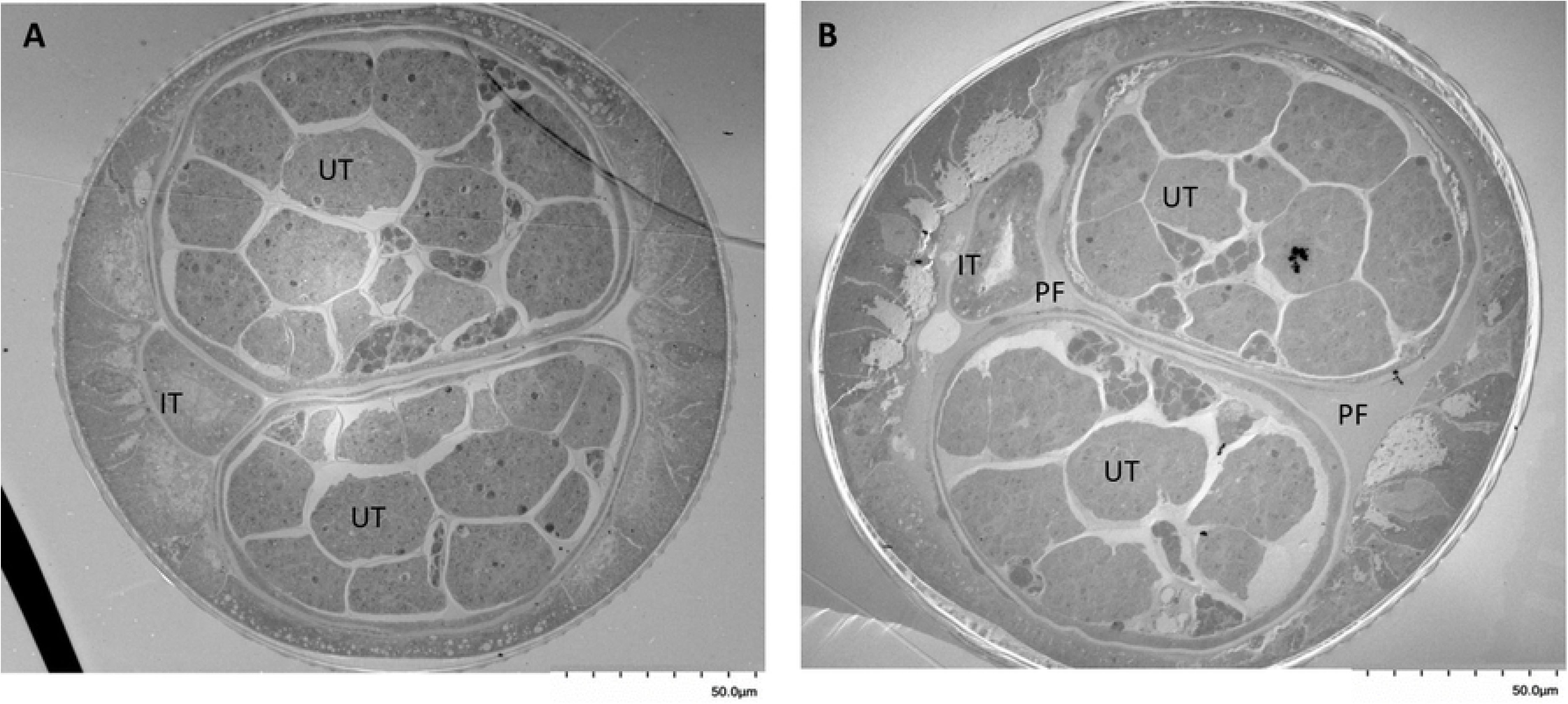
**Intestinal tract of a *B. malayi* adult worm treated with *Bma-*LAD-2 siRNA loses pseudocoelomic fluid.** The cross-sectional images were captured by transmission electron microscopy (TEM) at 500x. The adult filaria treated with siRNA (A) has lost most of the pseudocoelomic fluid surrounding the intestinal and uterine tubes, and appears to have fluid within the intestinal lumen. In contrast, the untreated filaria (B) has a normal distribution of pseudocoelomic fluid in the spaces around the intestinal and uterine tracts. PF = pseudocoelomic fluid, IT = intestinal tube, UT = uterine tube.

### No detectable Bma-LAD-2 specific IgG or IgE in serum from filarial patients

A major concern when evaluating possible vaccine candidates for helminths is whether populations in endemic areas are IgE-sensitized to the candidate antigen [27]. Thus, we investigated whether serum from filaria-infected individuals contains IgE that recognizes Bma-LAD-2. A luciferase immunoprecipitation system assay was employed to detect antibody levels in the patient serum samples using a Bma-LAD-2-luciferase fusion protein. Patients were categorized as presenting with asymptomatic microfilaremia (n=13), chronic pathology (lymphedema) (n=9), or tropical pulmonary eosinophilia (n=8). Sera used in this experiment were obtained from patients prior to anthelmintic treatment. We also tested sera from individuals with no clinical or laboratory evidence of a filarial infection (n=5) as well as healthy sera from blood bank donors (n=5). As positive controls, we used affinity-purified polyclonal antibodies raised in rabbits immunized with recombinant Bma-LAD-2 as well as the rabbit anti-sera. Naïve rabbit sera served as a negative control.

We found that serum samples from patients with lymphatic filariasis had no detectable pre-existing IgG (Fig 9A) or IgE (Fig 9B) against Bma-LAD-2 fusion protein. As expected, there was recognition by the polyclonal antibodies and anti-sera against the fusion protein. This indicated that our fusion protein exhibited the proper conformation and thus the absence of signal in filarial patient samples was due to absence of Bma-LAD-2 specific IgG or IgE.

**Fig 9.**
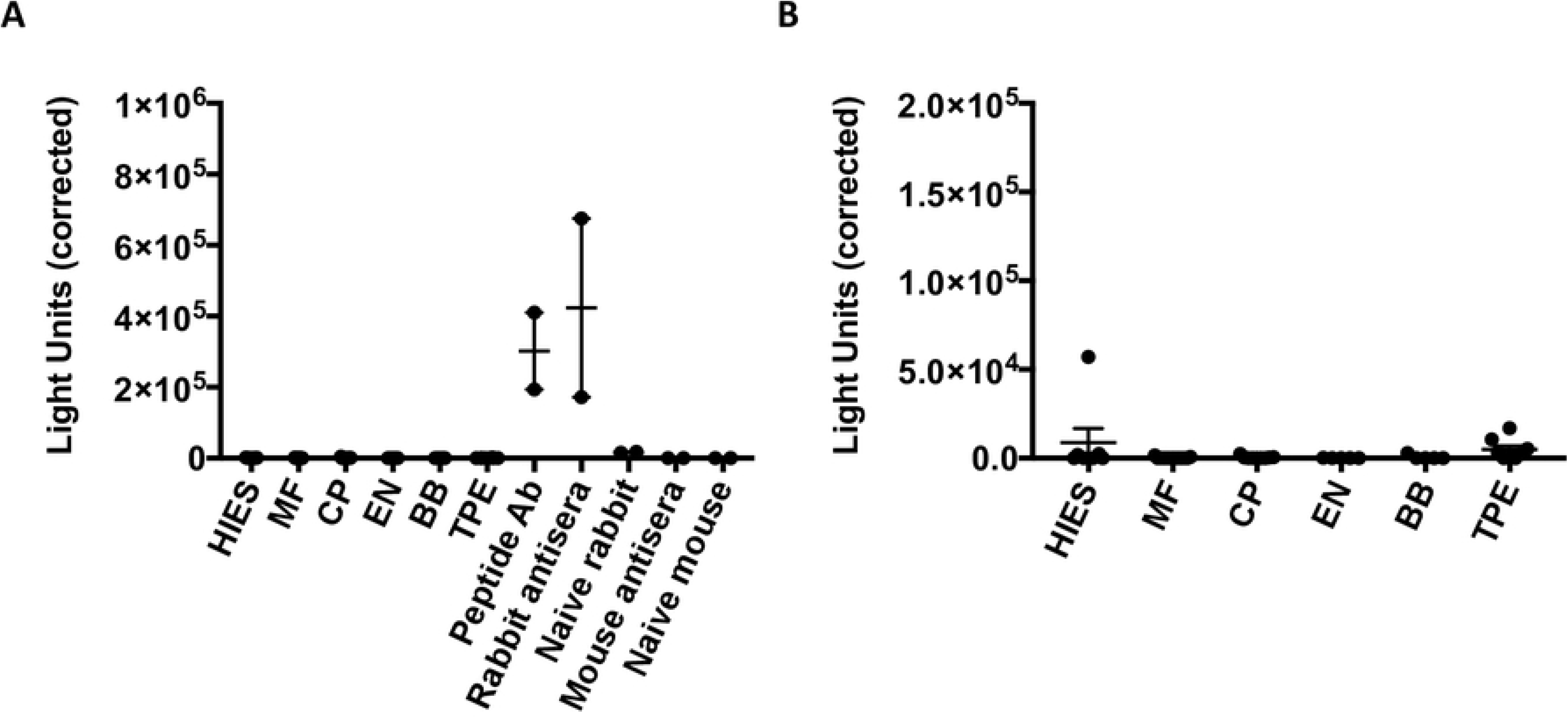
**Filarial patient serum does not contain detectable IgG or IgE against *Bma-*LAD-2.** Serum from filaria-infected individuals was incubated with a *Bma-*LAD-2-luciferase fusion protein. There was no detectable (A) IgG and (B) IgE in the patient serum as measured by the LIPS assay. As a positive control for IgG binding, *Bma-*LAD-2 rabbit polyclonal antibodies recognized the fusion protein. HIES=hyper IgE syndrome, MF=microfilaremic, CP=chronic pathology, EN=endemic normal, BB=blood bank donors, TPE=tropical pulmonary eosinophilic, Peptide Ab=*Bma-*LAD-2 rabbit polyclonal antibodies, Mouse antisera=serum from *Litomosoides sigmodontis* vaccinated mice

### siRNA knockdown of other intestinal antigens of *B. malayi*

In this study, we also attempted to evaluate whether 7 other intestinal proteins were essential for adult *B. malayi* survival (Table 1). The proteins selected for investigation were annotated as adhesion molecules, proteases, protease inhibitors, or involved in glycosylation based on work from previous studies [22, 26]. We were able to successfully knockdown gene expression for 3 of these target proteins. This limited success with siRNA inhibition was not entirely unexpected given the reported variability and difficulty of performing RNA interference in parasitic nematodes [28, 29]. Of note, unlike knockdown of Bma-LAD-2, successful siRNA inhibition of Bma-serpin (a protease inhibitor) and Bma-reprolysin (a protease) did not cause any appreciable phenotypic changes in adult *B. malayi* worms. siRNA inhibition of Bma-shtk, also a predicted protease, caused only minimal decreases in adult worm motility and metabolism.

**Table 1.**
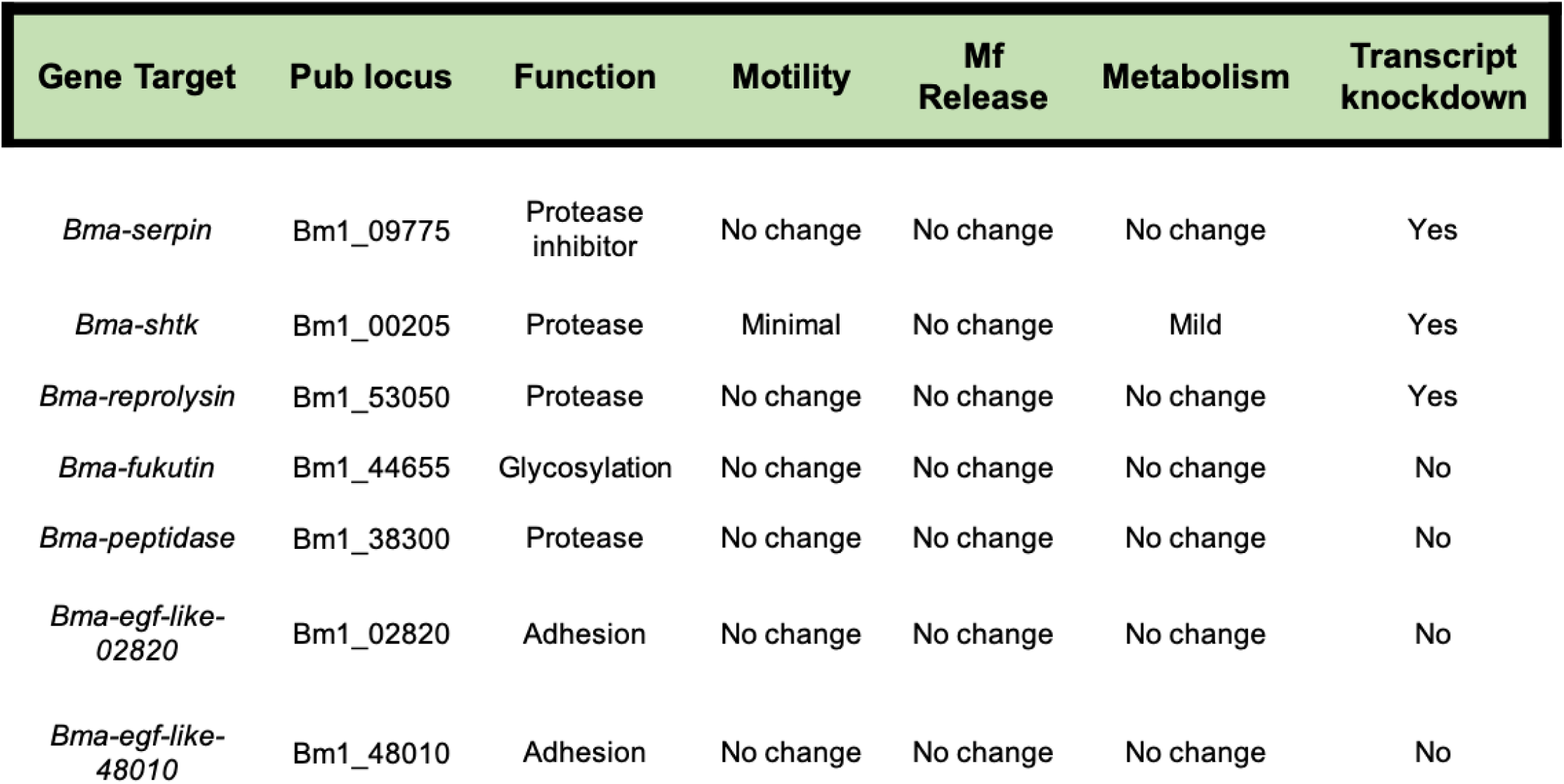
siRNA experiments targeting other intestinal proteins in *B. malayi* adult female worms.

| Gene Target | Pub locus | Function | Motility | Mf Release | Metabolism | Transcript knockdown |
| --- | --- | --- | --- | --- | --- | --- |
| *Bma-serpin* | Bm1_09775 | Protease inhibitor | No change | No change | No change | Yes |
| *Bma-shtk* | Bm1_00205 | Protease | Minimal | No change | Mild | Yes |
| *Bma-reprolysin* | Bm1_53050 | Protease | No change | No change | No change | Yes |
| *Bma-fukutin* | Bm1_44655 | Glycosylation | No change | No change | No change | No |
| *Bma-peptidase* | Bm1_38300 | Protease | No change | No change | No change | No |
| *Bma-egf-like-02820* | Bm1_02820 | Adhesion | No change | No change | No change | No |
| *Bma-egf-like-48010* | Bm1_48010 | Adhesion | No change | No change | No change | No |

## Methods

### Parasites and culture

*B. malayi* female adults were obtained from the NIH/NIAID Filariasis Research Reagent Resource Center (FR3) and TRS Laboratories in Athens, Georgia, USA. Before siRNA inhibition, adult worms were incubated in Dulbecco’s Modified Eagle’s Medium (Corning Cellgro) supplemented with 10% heat-inactivated fetal bovine serum (Atlanta Biologicals), 100 units/mL of penicillin, 100 ug/mL of streptomycin, and 1% L-glutamine (Sigma) for 24 hrs at 37°C in 5% CO_2_. Infection studies conducted at FR3 and TRS received approval from their respective Animal Care and Use Committees. Protocol approval for receipt of filarial worms from FR3 and TRS for use at the Uniformed Services University of the Health Sciences (USUHS) was granted by the USUHS Animal Care and Use Committee.

### Phylogenetic tree analysis

We investigated the degree of relatedness between helminth orthologs and Bma-LAD-2. As an outlier group for the phylogenetic tree, we included dogs, humans, and cats, which were expected to have a significantly distant relation to Bma-LAD-2 given the low predicted sequence homology. The tree was constructed based on the likelihood estimation method for the LG model using aligned sequences by MUtiple Sequence Comparison by Log-Expectation (MUSCLE)

Orthologs from nematode species were selected using a BLAST query of the WormBase Parasite database [30] against the Bma-LAD-2 protein sequence (WBGene00227085). The following are the nematode species along with the accession codes for each ortholog identified: *Brugia timori* (BTMF_0000455001), *Wuchereria bancrofti* (maker-PairedContig_4689-snap-gene-5.23)*, Brugia pahangi* (BPAG_0001424601)*, Loa loa* (LOAG_18710)*, Dirofilaria immitis* (nDi.2.2.2.t02266),, *Haemonchus contortus* (HCON_00104790)*, Necator americanus* (NECAME_12511), Onchocerca volvulus (Ovo-lad-2), *Caenorhabditis elegans* (lad-2), *Acanthocheilonema viteae* (nAv.1.0.1.t02543-RA), and *Ancylostoma duedonale* (ANCDUO_13310).

Orthologs from select mammals were identified using a BLAST query of the National Center of Biotechnology Information (NCBI) databases for the Bm-UGT peptide sequence. The following are the orthologs selected for analyses: *Homo sapiens* (NP_001153805.1), *Canis lupus familiaris* (XP_005640833.2), and *Felis catus* (XP_023103545.1).

### Structural analysis of Bma-LAD-2

Molecular model of the monomer and dimer of the extracellular domain of Bma-LAD-2 (residues 19-1120) was generated based on available structures/oligomerization modes of homologous proteins. The Ig1-Ig4 region was modeled using the structure of neurofascin, a member of the L1 family of neural cell adhesion molecules (sequence identity of 31%, PDB code: 3P3Y [24] and Ig5-6 and FN1-3 based upon the structures of contactin-3-6 (CNTN3-6), a group of glycophosphatidylinositol-anchored cell adhesion molecules (sequence identity of 28%, PDB code: 5I99, [31] and FN4-5 based upon the structure of a fragment encompassing the first four FN domains of the leucocyte common antigen-related protein (LAR), a post-synaptic type I transmembrane receptor protein (sequence identity 27%, PDB code: 6TPW) [32]. The dimer was assembled using the structure of neurofascin (PDB codes: 3P3Y and 3P40, [24] that assembles into symmetrical dimers in the crystal. The figure was generated using the PyMOL Molecular Graphics System, Version 2.0 Schrödinger, LLC (https://pymol.org/2/).

### siRNA for RNAi

BLOCK-iT^TM^ RNAi Designer was employed for selecting siRNA duplexes of candidate genes for gene silencing activity and specificity. The siRNA sequence with the greatest probability of success was selected for each target, and for some targets multiple sequences were selected to improve knockdown success. Life Technologies synthesized the target-specific siRNAs for Bma-LAD-2, Bma-Fukutin, Bma-ShTK, and Bma-Serpin and purified the complexes by standard desalting methods. Target-specific siRNAs for Bma-EGF-like-02820, Bma-EGF-like-48010, Bma-Peptidase, and Bma-Reprolysin were obtained through Dharmacon. The 5’-3’ siRNA sequences used in this experiment are as follows:

Bma-LAD-2 siRNA 1

sense: 5’ GCAAGUACUACCAUACUAUdTdT 3’
antisense: 5’ AUAAGUUGGAAUUCGUUGCdTdT 3’
Bma-LAD-2 siRNA 2

sense: 5’ GCGCAUAUCGCAAGUAAAUdTdT 3’
antisense: 5’ AUUUACUUGCGAUAUGCGCdTdT 3’
Bma-LAD-2 siRNA 3

sense: 5’ GCGAAUAGUCGAUACCUAAdTdT 3’
antisense: 5’ UUAGGUAUCGACUAUUCGCdTdT 3’
Bma*-*Fukutin siRNA 1

sense: 5’ CCACCCATTTCGCAGATTT 3’
antisense: 5’ AAAUCUGCGAAAUGGGUGGdTdT 3’
Bma*-*Fukutin siRNA 2

sense: 5’ GGAGCGAGAGTGAATGGAAdTdT 3’
antisense: 5’ UUCCAUUCACUCUCGCUCCdTdT 3’
Bma*-*Fukutin siRNA 3

sense: 5’ GCTAACGTTGCAAATTATTdTdT 3’
antisense: 5’ AAUAAUUUGCAACGUUAGCdTdT 3’
Bma*-*ShTK siRNA 1

sense: 5’ GCGCCTTCTACAGCAGTAAdTdT 3’
antisense: 5’ GCGCCUUCUACAGCAGUAAdTdT 3’
Bma*-*ShTK siRNA 2

sense: 5’ GGUGGUAUGAAUAGCAUAAdTdT 3’
antisense: 5’ UUAUGCUAUUCAUACCACCdTdT 3’
Bma*-*ShTK siRNA 3

sense: 5’ GCUAAAGAACUAUGCGCUAdTdT 3’
antisense: 5’ UAGCGCAUAGUUCUUUAGCdTdT 3’
Bma*-*Serpin siRNA

sense: 5’ GGAUUUCGAGUGAGACAAAdTdT 3’
antisense: 5’ UUUGUCUCACUCGAAAUCCdTdT 3’
Bma*-*EGF-like-02820 siRNA

sense: 5’ GUAUCGAGGGCAAGGGAAAdTdT 3’
antisense: 5’ UUUCCCUUGCCCUCGAUACdTdT 3’
Bma*-*EGF-like-48010 siRNA

sense: 5’ GCAACAAAUGCAAGAAUAAdTdT 3’
antisense: 5’ UUAUUCUUGCAUUUGUUGCdTdT 3’
Bma*-*Peptidase siRNA 1

sense: 5’ AGGAAAGGUUGUUAGGAUAdTdT 3’
antisense: 5’ UAUCCUAACAACCUUUCCUdTdT 3’
Bma*-*Reprolysin siRNA 3

sense: 5’ GGAUAAUGUGAAAGGAAUAdTdT 3’
antisense: 5’ UAUUCCUUUCACAUUAUCCdTdT 3’

### Assessment of siRNA uptake by fluorescence microscopy

Adult female worms were incubated with 5 μM of 5’ cy3-labeled *Bma-lad-2* siRNA 1 (Sigma Aldrich) for 24 hrs to evaluate uptake of siRNA into intestinal tract epithelial cells. Adult female worms were cultured in media alone as a negative control. As a counterstain, samples were treated with 10 μg/mL of 4′,6-Diamidino-2-phenylindole dihydrochloride (DAPI, Sigma-Aldrich) in PBS. Images were obtained with a Nikon Eclipse E600 fluorescent microscope and converged by NIS-Elements software.

### siRNA treatment of *B. malayi*

For siRNA inhibition of the target gene expression, we soaked *B. malayi* adult female worms in culture media with siRNA slightly modifying a previously established protocol [33]. For each timepoint, we incubated 5 adult female worms for 24 hrs in an equal mixture of the siRNAs (Bma-LAD-2, Bma-Fukutin, Bma-ShTK, Bma-Serpin) or one siRNA (Bma-EGF-like-02820, Bma-EGF-like-48010, Bma-Peptidase, Bma-Reprolysin) at a total concentration of 5 μM in 850 μL of media in a 5000 MWCO Pur-A-Lyzer^TM^ dialysis tube (Sigma-Aldrich). Previous studies have shown that this concentration of siRNA sufficiently silences gene expression [33–36]. We placed the dialysis tubes in a beaker with 500 mL of culture media at 37°C in 5% CO_2_. For the control groups, 5 adult female worms were incubated alone in media or with scrambled siRNA (5 μM) under conditions similar to the target siRNA-treated group. The worms were extracted after the 24-hr incubation and placed individually into 1 mL of culture media. Initially, worms were evaluated 1 day post-incubation for transcript knockdown, worm motility, MTT reduction, and microfilariae release. For Bma-LAD-2, we conducted additional experiments to evaluate the worms at 6 days post-siRNA incubation.

### Evaluation of worm motility

Motility was evaluated based on the following scale 4 = active movement, 3 = modest reduction in movement, 2 = severe reduction in movement, 1 = twitching, and 0 = no movement. A blinded observer rated worm motility for each group under a dissecting microscope.

### Measuring MTT reduction

Metabolic function was evaluated using a (4,5-dimethylthiazol-2-yl)-2,5-diphenyltetrazolium bromide (MTT) assay from Sigma [37]. For each group per timepoint, 2 worms were treated with 0.5 mg/mL of MTT in 0.5 mL of PBS for 30 minutes at 37°C in 5% CO_2_. Each worm was then transferred into a well containing 200 μL of DMSO of a 96-well plate at room temperature for 1 hr. Quantification of MTT reduction was measured based on absorbance relative to a DMSO blank at 570 nm with a Synergy HTX multi-mode plate reader (BioTek).

### Quantifying microfilaria (Mf) release

Prior to quantifying Mf release, we incubated the adult worms in 1 mL of fresh media for 24 hrs. The adult filariae were then removed for evaluation by the MTT reduction assay and RT-qPCR. The Mf were enumerated under a light microscope at high magnification in the wells containing expended media.

### RNA extraction and analysis of RNA levels by RT-qPCR

Adult *B. malayi* female worms were treated with TRIzol (Thermo Fischer Scientific) and subjected to three freeze/thaw cycles. Adult filariae were then placed in Matrix D lysis tubes (MP Biomedicals) and homogenized by a FastPrep^TM^-24 Biopulverizer (MP Biomedicals) for 7 minutes at 6 m/s. We added chloroform to the homogenate and phase separated the mixtures in Phase Lock Gel tubes (5Prime) at 11,900 g for 15 minutes at 4°C. After the top layer (aqueous phase) was collected, we precipitated the RNA using cold isopropanol and then pelleted it at 12,000 g for 1 hr. The RNA pellet was washed twice using cold 75% ethanol. We resuspended the RNA in nuclease-free water and quantified the sample concentrations using a NanoDrop 1000 (Thermo Fischer Scientific). Using Superscript IV (Thermo Fischer Scientific), we synthesized cDNA from mRNA per the manufacturer’s protocol. We quantified target gene and *B. malayi* house-keeping gene *gapdh* expression levels in duplicate 20 μL reactions using 1 μL of 20X TaqMan^TM^ gene expression assay (Thermo Fischer), 1 μL of cDNA, and 18 μL of TaqMan^TM^ gene expression master mix (Applied Biosystems). We employed the following PCR conditions: 2 min at 50 °C, 10 min at 95 °C, 40 cycles of 15 sec at 95 °C, and 1 min at 60 °C cycle of 50 °C with a 7500 Real-Time PCR System (Applied Biosystems). The following Taqman primer and internal probes were used:

Bma-gapdh:

Forward primer: 5’ TTGATCTCACTTGCCGACTC 3’
Reverse primer: 5’ TGGTCTTCGGTGTATTCCAA 3’
Internal probe: 5’ CAGCTAATGGACCGATGAAGGGGA 3’
Bma-lad-2:

Forward primer: 5’ GTGATCCACGGCTTACGATT 3’
Reverse primer: 5’ CAGGCACATCAAGCACAGTT 3’
Internal probe: 5’ TGCTCGTGGCTTTCATTCAGGA 3’
Bma-futkin:

Forward primer: 5’ AGGTTATTTCATGTGCCCTGC 3’
Reverse primer: 5’ ATTCCATTCACTCTCGCTCCA 3’
Internal probe: 5’ AGGCGGATTACGGTAATTGGCGAGT 3’
Bma-shtk:

Forward primer: 5’ TGCACTGATCCAATGGCAAA 3’
Reverse primer: 5’ GTTACTGCTGTAGAAGGCGC 3’
Internal probe: 5’ TGCGCCAAAACATGTGGATTTTGCGG 3’
Bma-serpin:

Forward primer: 5’ ACGTGCGCAGTTAGACTTTG 3’
Reverse primer: 5’ GCCTCTGCGATATAAGCCAA 3’
Internal probe: 5’ GCGGACGGTGAAACGAAGCAGCA 3’
Bma-egf-like-02820:

Forward primer: 5’ GCTTACACGGTGGCAGAAAA 3’
Reverse primer: 5’ AAGCCACCTATCTGCTCTCC 3’
Internal probe: 5’ TCGAGGGCAAGGGAAAACTGGAA 3’
Bma-egf-like-48010:

Forward primer: 5’ ACCTGGCTTCATGGGAGAAA 3’
Reverse primer: 5’ CTTCACCACAGTCGCAAACA 3’
Internal probe: 5’ TGCTGCCGGTCTTATGGGCG 3’
Bma-peptidase:

Forward primer: 5’ CAGCCATTATTGGCCAGGAC 3’
Reverse primer: 5’ AAATGAAGTGGTGCCGCATT 3’
Internal probe: 5’ AGCCTTCCAACTTGGTTCATCCCAACA 3’
Bma-reprolysin:

Forward primer: 5’ TGGAACACAGTGATCAGGCT 3’
Reverse primer: 5’ AACGGCATTCCACTTATCG 3’
Internal probe: 5’ CCCATTTCGTGTGCAATAGTTGCAGCA 3’

### Generation of anti-Bma-LAD-2 polyclonal antibodies

For the immunoblot analysis and LIPS assay, polyclonal anti-Bma-LAD-2 peptide antibodies were generated by Genscript. Rabbits were immunized with Bma-LAD-2 peptide sequences conjugated to keyhole limpet hemocyanin (KLH). The peptide sequences used are as follows: CYEKDEHLIAEGRPN, DSTGSKLAKTVKIDC, and CGQIANFDPYGRKMS. To facilitate binding to KLH, cysteines were added at either the N- or C-terminus of the peptides.

### Immunoblot analysis of Bma-LAD-2

Prior to Western blot analysis, we incubated *B. malayi* adult female worms in 5 μM of Bma-LAD-2 siRNA for 24 hrs followed by an additional 24 hr culture in fresh media. The adult filariae were transferred into Matrix D lysis tubes (MP Biomedicals) with PBS (pH 7.4) and 4 μL of Halt^TM^ Protease Inhibitor Cocktail (Thermo Scientific) and then homogenized using a FastPrep^TM^-24 Biopulverizer (MP Biomedicals) for 3 min at 4 m/s. Protein concentration was quantified by Bradford protein assay (Bio-Rad). Protein lysate (10 μg) was separated on 10% Bis-Tris NuPAGE gel (Invitrogen) and then transferred onto a 0.2 μm nitrocellulose filter paper (Bio-Rad). The filter paper was blocked overnight in 5% bovine serum albumin (BSA) in tris-buffered saline with 0.1% Tween 20 (TBS-T). After the overnight blocking, the membrane was incubated with 1:7000 polyclonal anti-Bma-LAD-2 peptide antibodies (Genscript) and 1:1000 rabbit anti-β actin antibodies (Abcam) for 1 hr. The membrane was washed three times with TBS-T for 15 min. Horseradish peroxidase conjugated goat anti-rabbit IgG antibody was incubated for 1 hr with the filter paper at a dilution of 1:2000. After washing again with TBS-T, the membrane was developed with Chemiluminescent reagent, SuperSignal^TM^ West Pico PLUS (Thermo Scientific).

### Transmission electron microscopy

*B. malayi* female worms (3) were treated with Bma-LAD-2 siRNA for 24 hrs and then cultured for an additional 48 hrs. An equal number of adult female worms were incubated in media alone for same amount of time. Both groups of filariae were processed for imaging by electron microscopy. This whole process was repeated on two separate occasions. For morphological evaluation, the female filariae were first washed in PBS (pH 7.4) and then fixed with 2.5% paraformaldehyde, 1% glutaraldehyde in 0.12 M Millong’s phosphate buffer (pH 7.4) overnight at room temperature. Following this step, the samples were post-fixed with 1% osmium tetroxide in 0.12 M Millong’s phosphate buffer (pH 7.4) for 100 min and then fixed en bloc with 2% aqueous uranyl acetate for 90 min. The samples were dehydrated in graded ethanol solutions (75% to 100%) for 10 min each. The worms were embedded in low viscosity epoxy resin (modified Spurr’s recipe) and then dried at 70°C overnight. Ultrathin sections (75 nm) were cut on a Reichert Ultracut E Ultramicrotome and then stained with 0.2% lead citrate. Reagents used were obtained from Electron Microscopy Services. Samples were visualized using a Hitachi HT7700 Transmission Electron Microscope at an accelerating voltage of 80 kV.

### Ruc-antigen fusion protein

The Bma-LAD-2-*Renilla reniformis* luciferase (Ruc) construct was inserted into a pREN2 vector by Genscript. The predicted signal sequence was removed prior to gene synthesis. The vector was cloned into TOP10 cells (Thermofischer) and amplified on agarose plates with kanamycin 50 μg/mL. Plasmid DNA was isolated and purified using a Miniprep kit (Qiagen) per the manufacturer’s guidelines. 293F cells (Thermofischer Scientific) were transfected with 30 μg of Bma-LAD-2 plasmid at a concentration of 1 x 10^6^ cells per mL. The 293F cells were collected 72 hrs later and homogenized. The lysate was stored at -80°C for later use.

### Luciferase immunoprecipitation system (LIPS)

We employed the LIPS assay to measure antibody titers in serum from *W. bancrofti* infected patients [38–40]. In a 96-well polypropylene plate, serum was diluted to 1:100 for IgG and 1:10 for IgE in 50 μL of LIPS master mix (20 mM Tris pH 7.5, 150 mM NaCl, 5mM MgCl2, 1% Triton X-100) with PBS-T added to bring the volume to 100 μL. We added 1 x 10^6^ light units (LU) of the LAD-2-Ruc fusion protein to the mixture and incubated it at room temperature for 10 min. Purification of the antigen-antibody complex involved adding 5 μL of a 50% suspension of Ultralink protein A/G (Pierce) or Ultralink anti-human IgE beads in PBS to a 96-well filter plate (Milipore) and then applying a vacuum. The serum mixture was then added and allowed to incubate for 15 minutes at room temperature. A vacuum was applied to the filter plate leaving only antigen-antibody complexed bound to beads in the wells. The samples were washed with 200 μL of LIPS master mix twice and with PBS once. Using Bethold LB 960 Centro microplate luminometer, emitted LUs were measured after addition of 50 μL of coelenterzine solution (Promega) to each sample well. The serum samples were run in duplicate, and the calculated LU was the emitted LU for only the LAD-2 fusion protein subtracted from the emitted LU for each sample.

### Serum Samples

Serum samples used in this study were obtained under Institutional Review Board (IRB)-approved protocols from the Department of Transfusion Medicine (Clinical Center, National Institutes of Health, Bethesda, MD) and from the Laboratory of Laboratory of Parasitic Diseases (NIAID, National Institutes of Health, Bethesda, MD). All donors provided written approved consent.

### Statistical analysis

The siRNA experiments for Bma-LAD-2 were repeated twice under the same conditions. All other siRNA experiments were only performed once. The worm motility and Mf release data was analyzed by one-way analysis of variance (ANOVA) using the statistical package in PRISM 7.0. Validity of the one-way ANOVA was verified by performing individual comparisons of mean values using Tukey’s multiple comparisons test. For the gene expression and MTT reduction data, a T-test was used to determine significance. The p values for each experimental and control group was designated as follows: * for p values <0.05, ** for p values <0.01, and *** for p values <0.001.

## Discussion

In this study, we sought to identify intestinal tract antigens of adult filarial nematodes that could potentially serve as therapeutic or vaccine targets. Even though Bma-LAD-2 is expressed in all lifecycle stages, we hypothesize that disruption of tight junctions formed within the intestinal tract of filariae will have a selective effect on the adult worm stages as microfilariae lack an intestinal tract [41]. Knockdown of the tight junction protein Bma-LAD-2 caused rapid reductions in worm motility, metabolism, and microfilaria release, which led to worm death. Electron microscopy demonstrated a loss of pseudocoelomic fluid, revealing that disruption of the tight junctions between epithelial cells of the filarial intestinal tract could be a novel method to rapidly kill these worms by causing rapid dissolution of their hydrostatic skeleton.

Bma-LAD-2 is an immunoglobulin (Ig) intermediate-set (I-set) domain containing protein and, therefore, belongs to the functionally diverse Ig superfamily (IgSF). The Ig domain is the basic structural unit of the superfamily and consists of two sandwiched antiparallel beta sheets [42]. Proteins in the IgSF are classified based on the structure of their Ig domain and given a set designation of variable (V), constant 1 (C1), constant 2 (C2), or intermediate (I) [43]. Ig I-set domains are similar to V-set domains but have a shorter distance between the invariant cysteine residues. Intropro analysis predicts that Bma-LAD-2 has 6 Ig domains spanning from amino acid position 24 to 602.

Based on homology to its ortholog in *C. elegans* (LAD-2, L1 adhesion), Bma-LAD-2 is predicted to be an L1 cell adhesion molecule (L1CAM). L1CAMs are single transmembrane proteins that can participate in homophilic and heterophilic interactions [44]. The cytoplasmic tail of L1CAMs has multiple consensus-binding sites which allow for interaction with various cytoskeleton linkers proteins such as ankyrin, spectrin, and ERM [45, 46]. Furthermore, the cytoplasmic tail of L1CAMs has phosphorylation sites indicating a possible role in signal transduction [44, 47, 48].

Interestingly in *C. elegans,* LAD-2 is critical in guiding axon migration and does not appear to be critical for the establishment or maintenance of the intestinal epithelium [49, 50]. However, LAD-1, another L1CAM, has been shown to co-localize with apical junction molecules. In nematodes, cell adhesion molecules (CAMs) assemble to form two major types of apical junctions: the cadherin catenin complex (CCC) and the DLG-1/AJM-1 complex (DAC) [45, 50]. In *C. elegans*, it has been shown that that the CCC is not essential for cell adhesion [51]. This is surprising given the critical role of cadherins in cellular adhesion for most other eukaryotes. Researchers have suggested that LAD-1 may act as a redundant adhesion system thereby mitigating the loss of the CCC. Indeed, embryos expressing dominant-negative LAD-1 have altered cell morphology and position indicating a role in cellular adhesion [52–54]. In filarial nematodes, no such redundancy appears to exist as knockdown of Bma-LAD-2 resulted in dramatic phenotypic change.

Evidence also indicates that L1CAMs play an integral role in cell-to-cell contact signaling in the epithelial cells. In fact, loss of L1CAM signal can arrest cell proliferation and potentially induce apoptosis [55, 56]. This would explain the ablation of microvilli in the intestine of target specific siRNA-treated worms as well as the unraveling of the mitochondrial cristae. In addition, the loss of adhesion molecules may have hindered the ability of the apical junction to prevent diffusion of the internal pseudocoelomic fluid into the intestinal lumen. Loss of this fluid would have an adverse effect on worm vitality. The pseudocoelomic fluid generates a positive pressure within the pseudocoelom creating a hydrostatic skeleton thought to be necessary for maintaining cuticle rigidity. This allows the longitudinal musculature of the worm to contract against the cuticle creating the wave movement necessary for locomotion [41]. In addition, it believed that the fluid serves as lubricant for the tissues during this motile process as well as a medium for nutrient exchange and cellular signaling [57]. After an extensive search of the current literature, this loss of pseudocoelomic fluid appears to be a unique finding, and it most likely played a significant role in establishing the phenotype seen with Bma-LAD-2 knockdown.

In addition to disrupted anatomy, changes in the intestinal tract may have contributed to worm death by disrupting nutrient intake. While studies have shown that *Brugia* worms can absorb some nutrients through their cuticle, [41, 58], a previous study of the rat filarial *Litomoisoides sigmoiditis* showed the presence of red blood cells in the filarial gut which implied that adult filariae actively feed [59]. Another study demonstrated that heartworms are able to ingest serum [60]. In addition, the proteomic analysis of different filarial tissue structures performed by our lab revealed that the filarial intestine is enriched in transporters, drug metabolizing enzymes, proteases, protease inhibitors, and adhesion molecules [22]. These findings suggest that the gut is used for not only nutrient digestion and uptake but also waste metabolism and disposal; functions essential in any living organism.

The loss of Bma-LAD-2 results in a different phenotype than what has been observed when the ortholog of this protein is knocked out in *C. elegans*. This could be due to a number of reasons. *C. elegans* constantly use their intestine for digestion and waste disposal, emptying their intestinal contents every 45 seconds [58, 61, 62]. Additionally, *C elegans* are unable to absorb nutrients across their thick cuticle, which leaves the intestine as the only means of nutrient absorption. Evolutionarily, it is reasonable that the systems in the intestine of *C. elegans* are redundant as failure in one could result in worm death. Indeed, we see evidence of this redundancy by the fact that knocking out the CCC does not dramatically disrupt cell adhesion in *C. elegans* [51]. In contrast, intestinal feeding by adult filariae may be more inconsistent as the parasites can absorb at least some nutrients through their cuticle [58, 62]. It is quite possible that helminths only use their intestine to digest essential proteins and macromolecules too large to be absorbed by the cuticle. Consequently, there may have been less evolutionary pressure to develop redundant systems in the intestine. Finally, *C elegans* is a free-living nematode, and therefore, has different digestive requirements than a parasitic nematode such as *Brugia*. This difference is no more apparent than with the number of protein-coding genes. *C elegans* has 19,762 protein-coded genes compared to the predicted ∼11,500 of *B. malayi* [26, 63].

When developing a helminth vaccine, there is a risk that the antigen could induce an allergic reaction in individuals previously exposed to filariae [21]. This has been a major impediment to the development of effective helminth vaccines. There is evidence to suggest that helminth intestinal proteins act as “hidden antigens,” which are proteins not exposed to the immune system during natural infection and thus would not elicit an IgE-response [16, 17, 21, 64, 65]. Furthermore, because these proteins are hidden from the immune system, there is little evolutionary pressure to develop mechanisms that enable these proteins to evade the immune system. This may leave these proteins vulnerable to attack by the host defenses [65]. A key limitation to development of vaccines against filarial nematodes is the possibility that endemic populations may be IgE-sensitized against the antigen and thus experience allergic reactions when vaccinated. In this study, we demonstrated that people infected with lymphatic filariasis do not appear to have detectable IgE antibodies against Bma-LAD-2, suggesting that this antigen may be safe to use as a vaccine. While this result is promising, more studies need to be performed using a larger sample size of endemic people to confirm these results.

In this study, we also evaluated 7 other *Brugia* intestinal proteins. Successful transcript knockdown was achieved for 3 of the targets. We attribute this failure rate to the well-documented difficulty of performing RNA interference in helminths [28, 29]. None of the other proteins that were successfully knocked down (Bma-Serpin, Bma-ShTK, and Bma-Reprolysin) resulted in substantial changes to worm viability or fecundity. It is expected that not all intestinal proteins are essential for adult filaria survival. Interestingly, inhibiting expression of Bma-Reprolysin, a putative protease, did not result in a noticeable phenotype. This may be due to the fact that there are multiple proteases present in the gut and that knockdown of more than one is necessary to affect worm survival.

Finally, we suspect that any therapeutics or vaccines developed against Bma-LAD-2 would be effective against *W. bancrofti* and *B. timori as well as B. malayi* due to high overall relatedness between the filaria species. In addition, Bma-LAD-2 shares a high level of homology (75%) with other filarial species. Therefore, a therapeutic or vaccine developed against this protein may be protective against other filarial infections.

In conclusion, we demonstrated knockdown of Bma-LAD-2 expression in adult *B. malayi* by siRNA inhibition. This resulted in ablation of microvilli, shortened tight junctions, unraveling of the mitochondrial cristae, and loss of pseudocoelmoic fluid as visualized by TEM. We believe that these structural changes in the intestinal epithelium led to deceased worm motility, metabolism, and Mf release in worms treated with Bma-LAD-2 siRNA. Therefore, we conclude that Bma-LAD-2 is an essential protein for adult worm survival. The lack of Bma-LAD-2-specific IgE suggests that this antigen would be a safe to use in a vaccine administered in endemic areas. In future studies we plan to evaluate Bma-LAD-2 as a vaccine candidate in animal models and as a potential therapeutic target for development of novel medications that specifically target adult filarial worms.

## Acknowledgements

We thank Belinda Jackson (Uniformed Services University) and Laura Kropp for their assistance with the worm motility scoring.

## Disclaimer

The opinions and assertions expressed herein are those of the author(s) and do not necessarily reflect the official policy or position of the Uniformed Services University, the Department of Defense, or the Department of Health and Human Services.

This work was prepared by military (AFF) and civilian (MEP, SB, TBN, EM) employees of the US Government as part of these individuals’ official duties and therefore is in the public domain and does not possess copyright protection *(public domain information may be freely distributed and copied; however, as a courtesy it is requested that the Uniformed Services University and the authors be given an appropriate acknowledgement)*.

## Data Availability

All relevant data are within the paper and its Supporting Information files.

## References

1. Ramaiah KD, Ottesen EA. Progress and impact of 13 years of the global programme to eliminate lymphatic filariasis on reducing the burden of filarial disease. PLoS Negl Trop Dis. 2014;8(11):e3319. doi: 10.1371/journal.pntd.0003319. PubMed PMID: 25412180; PubMed Central PMCID: PMC4239120.

2. Evaluation IfHMa. Global Burden of Disease Study 2019 Results Tool Seattle, Washington: University of Washington; 2021 [cited 2021 June 19]. Available from: http://ghdx.healthdata.org/gbd-results-tool.

3. Global Programme to Eliminate Lymphatic Filariasis: World Health Organization; 2016 [cited 2016 May 2]. Available from: http://www.who.int/lymphatic_filariasis/disease/en/.

4. WHO Team CoNTD. Ending the neglect to attain the Sustainable Development Goals: A road map for neglected tropical diseases 2021–2030. Overview. World Health Organization, 2020 Contract No.: WHO/UCN/NTD/2020.01.

5. Hoerauf A, Pfarr K, Mand S, Debrah AY, Specht S. Filariasis in Africa--treatment challenges and prospects. Clin Microbiol Infect. 2011;17(7):977–85. Epub 2011/07/05. doi: 10.1111/j.1469-0691.2011.03586.x. PubMed PMID: 21722251.

6. Rebollo MP, Bockarie MJ. Can Lymphatic Filariasis Be Eliminated by 2020? Trends Parasitol. 2017;33(2):83–92. doi: 10.1016/j.pt.2016.09.009. PubMed PMID: 27765440.

7. Bockarie MJ, Deb RM. Elimination of lymphatic filariasis: do we have the drugs to complete the job? Curr Opin Infect Dis. 2010;23(6):617–20. Epub 2010/09/18. doi: 10.1097/QCO.0b013e32833fdee5. PubMed PMID: 20847694.

8. Thomsen EK, Sanuku N, Baea M, Satofan S, Maki E, Lombore B, et al. Efficacy, Safety, and Pharmacokinetics of Coadministered Diethylcarbamazine, Albendazole, and Ivermectin for Treatment of Bancroftian Filariasis. Clin Infect Dis. 2016;62(3):334–41. Epub 2015/10/22. doi: 10.1093/cid/civ882. PubMed PMID: 26486704.

9. King CL, Suamani J, Sanuku N, Cheng YC, Satofan S, Mancuso B, et al. A Trial of a Triple-Drug Treatment for Lymphatic Filariasis. N Engl J Med. 2018;379(19):1801–10. Epub 2018/11/08. doi: 10.1056/NEJMoa1706854. PubMed PMID: 30403937; PubMed Central PMCID: PMCPMC6194477.

10. Twum-Danso NA. Loa loa encephalopathy temporally related to ivermectin administration reported from onchocerciasis mass treatment programs from 1989 to 2001: implications for the future. Filaria J. 2003;2 Suppl 1:S7. Epub 2004/02/21. doi: 10.1186/1475-2883-2-S1-S7. PubMed PMID: 14975064; PubMed Central PMCID: PMCPMC2147656.

11. Wanji S, Eyong EJ, Tendongfor N, Ngwa CJ, Esuka EN, Kengne-Ouafo AJ, et al. Ivermectin treatment of Loa loa hyper-microfilaraemic baboons (Papio anubis): Assessment of microfilarial load reduction, haematological and biochemical parameters and histopathological changes following treatment. PLoS Negl Trop Dis. 2017;11(7):e0005576. Epub 2017/07/08. doi: 10.1371/journal.pntd.0005576. PubMed PMID: 28686693; PubMed Central PMCID: PMCPMC5533442.

12. Albiez EJ, Newland HS, White AT, Kaiser A, Greene BM, Taylor HR, et al. Chemotherapy of onchocerciasis with high doses of diethylcarbamazine or a single dose of ivermectin: microfilaria levels and side effects. Trop Med Parasitol. 1988;39(1):19–24. Epub 1988/03/01. PubMed PMID: 3291074.

13. Awadzi K, Gilles HM. Diethylcarbamazine in the treatment of patients with onchocerciasis. Br J Clin Pharmacol. 1992;34(4):281–8. Epub 1992/10/01. PubMed PMID: 1457260; PubMed Central PMCID: PMCPMC1381407.

14. Bassetto CC, Silva BF, Newlands GF, Smith WD, Amarante AF. Protection of calves against Haemonchus placei and Haemonchus contortus after immunization with gut membrane proteins from H. contortus. Parasite Immunol. 2011;33(7):377–81. Epub 2011/05/04. doi: 10.1111/j.1365-3024.2011.01295.x. PubMed PMID: 21535018.

15. Loukas A, Bethony JM, Williamson AL, Goud GN, Mendez S, Zhan B, et al. Vaccination of dogs with a recombinant cysteine protease from the intestine of canine hookworms diminishes the fecundity and growth of worms. J Infect Dis. 2004;189(10):1952–61. doi: 10.1086/386346. PubMed PMID: 15122534.

16. Pearson MS, Bethony JM, Pickering DA, de Oliveira LM, Jariwala A, Santiago H, et al. An enzymatically inactivated hemoglobinase from Necator americanus induces neutralizing antibodies against multiple hookworm species and protects dogs against heterologous hookworm infection. FASEB J. 2009;23(9):3007–19. doi: 10.1096/fj.09-131433. PubMed PMID: 19380510; PubMed Central PMCID: PMC2735369.

17. Pearson MS, Pickering DA, Tribolet L, Cooper L, Mulvenna J, Oliveira LM, et al. Neutralizing antibodies to the hookworm hemoglobinase Na-APR-1: implications for a multivalent vaccine against hookworm infection and schistosomiasis. J Infect Dis. 2010;201(10):1561–9. Epub 2010/04/07. doi: 10.1086/651953. PubMed PMID: 20367477.

18. Zhan B, Liu S, Perally S, Xue J, Fujiwara R, Brophy P, et al. Biochemical characterization and vaccine potential of a heme-binding glutathione transferase from the adult hookworm Ancylostoma caninum. Infect Immun. 2005;73(10):6903–11. doi: 10.1128/IAI.73.10.6903-6911.2005. PubMed PMID: 16177370; PubMed Central PMCID: PMC1230892.

19. Zhan B, Perally S, Brophy PM, Xue J, Goud G, Liu S, et al. Molecular cloning, biochemical characterization, and partial protective immunity of the heme-binding glutathione S-transferases from the human hookworm Necator americanus. Infect Immun. 2010;78(4):1552–63. Epub 2010/02/11. doi: 10.1128/IAI.00848-09. PubMed PMID: 20145100; PubMed Central PMCID: PMCPMC2849424.

20. Hotez PJ, Beaumier CM, Gillespie PM, Strych U, Hayward T, Bottazzi ME. Advancing a vaccine to prevent hookworm disease and anemia. Vaccine. 2016;34(26):3001–5. Epub 2016/04/05. doi: 10.1016/j.vaccine.2016.03.078. PubMed PMID: 27040400.

21. Diemert DJ, Freire J, Valente V, Fraga CG, Talles F, Grahek S, et al. Safety and immunogenicity of the Na-GST-1 hookworm vaccine in Brazilian and American adults. PLoS Negl Trop Dis. 2017;11(5):e0005574. Epub 2017/05/04. doi: 10.1371/journal.pntd.0005574. PubMed PMID: 28464026; PubMed Central PMCID: PMCPMC5441635.

22. Morris CP, Bennuru S, Kropp LE, Zweben JA, Meng Z, Taylor RT, et al. A Proteomic Analysis of the Body Wall, Digestive Tract, and Reproductive Tract of Brugia malayi. PLoS Negl Trop Dis. 2015;9(9):e0004054. Epub 2015/09/15. doi: 10.1371/journal.pntd.0004054. PubMed PMID: 26367142; PubMed Central PMCID: PMCPMC4569401.

23. Flynn AF, Joyce MG, Taylor RT, Bennuru S, Lindrose AR, Sterling SL, et al. Intestinal UDP-glucuronosyltransferase as a potential target for the treatment and prevention of lymphatic filariasis. PLoS Negl Trop Dis. 2019;13(9):e0007687. Epub 2019/09/13. doi: 10.1371/journal.pntd.0007687. PubMed PMID: 31513587; PubMed Central PMCID: PMCPMC6742224.

24. Liu H, Focia PJ, He X. Homophilic adhesion mechanism of neurofascin, a member of the L1 family of neural cell adhesion molecules. J Biol Chem. 2011;286(1):797–805. Epub 2010/11/05. doi: 10.1074/jbc.M110.180281. PubMed PMID: 21047790; PubMed Central PMCID: PMCPMC3013039.

25. Choi YJ, Ghedin E, Berriman M, McQuillan J, Holroyd N, Mayhew GF, et al. A deep sequencing approach to comparatively analyze the transcriptome of lifecycle stages of the filarial worm, Brugia malayi. PLoS Negl Trop Dis. 2011;5(12):e1409. Epub 2011/12/20. doi: 10.1371/journal.pntd.0001409. PubMed PMID: 22180794; PubMed Central PMCID: PMCPMC3236722.

26. Bennuru S, Meng Z, Ribeiro JM, Semnani RT, Ghedin E, Chan K, et al. Stage-specific proteomic expression patterns of the human filarial parasite Brugia malayi and its endosymbiont Wolbachia. Proc Natl Acad Sci U S A. 2011;108(23):9649–54. Epub 2011/05/25. doi: 1011481108 [pii] 10.1073/pnas.1011481108. PubMed PMID: 21606368; PubMed Central PMCID: PMC3111283.

27. Diemert DJ, Pinto AG, Freire J, Jariwala A, Santiago H, Hamilton RG, et al. Generalized urticaria induced by the Na-ASP-2 hookworm vaccine: implications for the development of vaccines against helminths. J Allergy Clin Immunol. 2012;130(1):169–76 e6. doi: 10.1016/j.jaci.2012.04.027. PubMed PMID: 22633322.

28. Dalzell JJ, Warnock ND, McVeigh P, Marks NJ, Mousley A, Atkinson L, et al. Considering RNAi experimental design in parasitic helminths. Parasitology. 2012;139(5):589–604. doi: 10.1017/S0031182011001946. PubMed PMID: 22216952.

29. Ratnappan R, Vadnal J, Keaney M, Eleftherianos I, O’Halloran D, Hawdon JM. RNAi-mediated gene knockdown by microinjection in the model entomopathogenic nematode Heterorhabditis bacteriophora. Parasit Vectors. 2016;9:160. Epub 2016/03/20. doi: 10.1186/s13071-016-1442-4. PubMed PMID: 26993791; PubMed Central PMCID: PMCPMC4797128.

30. Altschul SF, Madden TL, Schaffer AA, Zhang J, Zhang Z, Miller W, et al. Gapped BLAST and PSI-BLAST: a new generation of protein database search programs. Nucleic Acids Res. 1997;25(17):3389–402. Epub 1997/09/01. PubMed PMID: 9254694; PubMed Central PMCID: PMCPMC146917.

31. Nikolaienko RM, Hammel M, Dubreuil V, Zalmai R, Hall DR, Mehzabeen N, et al. Structural Basis for Interactions Between Contactin Family Members and Protein-tyrosine Phosphatase Receptor Type G in Neural Tissues. J Biol Chem. 2016;291(41):21335–49. Epub 2016/08/20. doi: 10.1074/jbc.M116.742163. PubMed PMID: 27539848; PubMed Central PMCID: PMCPMC5076805.

32. Vilstrup J, Simonsen A, Birkefeldt T, Strandbygard D, Lyngso J, Pedersen JS, et al. Crystal and solution structures of fragments of the human leucocyte common antigen-related protein. Acta Crystallogr D Struct Biol. 2020;76(Pt 5):406–17. Epub 2020/05/02. doi: 10.1107/S2059798320003885. PubMed PMID: 32355037.

33. Aboobaker AA, Blaxter ML. Use of RNA interference to investigate gene function in the human filarial nematode parasite Brugia malayi. Mol Biochem Parasitol. 2003;129(1):41–51. Epub 2003/06/12. PubMed PMID: 12798505.

34. Kushwaha S, Singh PK, Shahab M, Pathak M, Bhattacharya SM. In vitro silencing of Brugia malayi trehalose-6-phosphate phosphatase impairs embryogenesis and in vivo development of infective larvae in jirds. PLoS Negl Trop Dis. 2012;6(8):e1770. Epub 2012/08/21. doi: 10.1371/journal.pntd.0001770. PubMed PMID: 22905273; PubMed Central PMCID: PMCPMC3419221.

35. Misra S, Gupta J, Misra-Bhattacharya S. RNA interference mediated knockdown of Brugia malayi UDP-Galactopyranose mutase severely affects parasite viability, embryogenesis and in vivo development of infective larvae. Parasit Vectors. 2017;10(1):34. Epub 2017/01/21. doi: 10.1186/s13071-017-1967-1. PubMed PMID: 28103957; PubMed Central PMCID: PMCPMC5244609.

36. Singh PK, Kushwaha S, Mohd S, Pathak M, Misra-Bhattacharya S. In vitro gene silencing of independent phosphoglycerate mutase (iPGM) in the filarial parasite Brugia malayi. Infect Dis Poverty. 2013;2(1):5. Epub 2013/07/16. doi: 10.1186/2049-9957-2-5. PubMed PMID: 23849829; PubMed Central PMCID: PMCPMC3707094.

37. Comley JC, Rees MJ, Turner CH, Jenkins DC. Colorimetric quantitation of filarial viability. Int J Parasitol. 1989;19(1):77–83. PubMed PMID: 2707965.

38. Burbelo PD, Goldman R, Mattson TL. A simplified immunoprecipitation method for quantitatively measuring antibody responses in clinical sera samples by using mammalian-produced Renilla luciferase-antigen fusion proteins. BMC Biotechnol. 2005;5:22. Epub 2005/08/20. doi: 10.1186/1472-6750-5-22. PubMed PMID: 16109166; PubMed Central PMCID: PMCPMC1208859.

39. Burbelo PD, Ramanathan R, Klion AD, Iadarola MJ, Nutman TB. Rapid, novel, specific, high-throughput assay for diagnosis of Loa loa infection. J Clin Microbiol. 2008;46(7):2298–304. Epub 2008/05/30. doi: 10.1128/JCM.00490-08. PubMed PMID: 18508942; PubMed Central PMCID: PMCPMC2446928.

40. Drame PM, Meng Z, Bennuru S, Herrick JA, Veenstra TD, Nutman TB. Identification and Validation of Loa loa Microfilaria-Specific Biomarkers: a Rational Design Approach Using Proteomics and Novel Immunoassays. MBio. 2016;7(1):e02132–15. Epub 2016/02/18. doi: 10.1128/mBio.02132-15. PubMed PMID: 26884435; PubMed Central PMCID: PMCPMC4791851.

41. Scott AL. Lymphatic-dwelling filariae. In: Nutman T, editor. Lymphatic Filariasis. London: Imperial College Press; 2000. p. 5–39.

42. Buck CA. Immunoglobulin superfamily: Structure, function and relationship to other receptor molecules. Seminars in Cell Biology. 1992;3(3):179–88. doi: http://dx.doi.org/10.1016/S1043-4682(10)80014-5.

43. Smith DK, Xue H. Sequence profiles of immunoglobulin and immunoglobulin-like domains. Journal of molecular biology. 1997;274(4):530–45. doi: 10.1006/jmbi.1997.1432. PubMed PMID: 9417933.

44. Kiefel H, Bondong S, Hazin J, Ridinger J, Schirmer U, Riedle S, et al. L1CAM: a major driver for tumor cell invasion and motility. Cell Adh Migr. 2012;6(4):374–84. Epub 2012/07/17. doi: 10.4161/cam.20832. PubMed PMID: 22796939; PubMed Central PMCID: PMCPMC3478260.

45. Hartsock A, Nelson WJ. Adherens and tight junctions: structure, function and connections to the actin cytoskeleton. Biochim Biophys Acta. 2008;1778(3):660–9. doi: 10.1016/j.bbamem.2007.07.012. PubMed PMID: 17854762; PubMed Central PMCID: PMC2682436.

46. Takahashi K, Nakanishi H, Miyahara M, Mandai K, Satoh K, Satoh A, et al. Nectin/PRR: An Immunoglobulin-like Cell Adhesion Molecule Recruited to Cadherin-based Adherens Junctions through Interaction with Afadin, a PDZ Domain–containing Protein. The Journal of Cell Biology. 1999;145(3):539–49. PubMed PMID: PMC2185068.

47. Chen L, Zhou S. "CRASH"ing with the worm: insights into L1CAM functions and mechanisms. Dev Dyn. 2010;239(5):1490–501. Epub 2010/03/13. doi: 10.1002/dvdy.22269. PubMed PMID: 20225255; PubMed Central PMCID: PMCPMC3428060.

48. Hoffmann M, Segbert C, Helbig G, Bossinger O. Intestinal tube formation in Caenorhabditis elegans requires vang-1 and egl-15 signaling. Dev Biol. 2010;339(2):268–79. Epub 2009/12/17. doi: 10.1016/j.ydbio.2009.12.002. PubMed PMID: 20004187.

49. Wang X, Zhang W, Cheever T, Schwarz V, Opperman K, Hutter H, et al. The C. elegans L1CAM homologue LAD-2 functions as a coreceptor in MAB-20/Sema2 mediated axon guidance. J Cell Biol. 2008;180(1):233–46. Epub 2008/01/16. doi: 10.1083/jcb.200704178. PubMed PMID: 18195110; PubMed Central PMCID: PMCPMC2213605.

50. Lynch AM, Hardin J. The assembly and maintenance of epithelial junctions in C. elegans. Front Biosci (Landmark Ed). 2009;14:1414–32. Epub 2009/03/11. PubMed PMID: 19273138; PubMed Central PMCID: PMCPMC2896272.

51. Costa M, Raich W, Agbunag C, Leung B, Hardin J, Priess JR. A putative catenin-cadherin system mediates morphogenesis of the Caenorhabditis elegans embryo. J Cell Biol. 1998;141(1):297–308. Epub 1998/05/16. PubMed PMID: 9531567; PubMed Central PMCID: PMCPMC2132712.

52. Wang X, Kweon J, Larson S, Chen L. A role for the C. elegans L1CAM homologue lad-1/sax-7 in maintaining tissue attachment. Dev Biol. 2005;284(2):273–91. Epub 2005/07/19. doi: 10.1016/j.ydbio.2005.05.020. PubMed PMID: 16023097.

53. Dubreuil RR. Functional links between membrane transport and the spectrin cytoskeleton. J Membr Biol. 2006;211(3):151–61. Epub 2006/11/09. doi: 10.1007/s00232-006-0863-y. PubMed PMID: 17091212.

54. Weiss EE, Kroemker M, Rudiger AH, Jockusch BM, Rudiger M. Vinculin is part of the cadherin-catenin junctional complex: complex formation between alpha-catenin and vinculin. J Cell Biol. 1998;141(3):755–64. Epub 1998/06/13. PubMed PMID: 9566974; PubMed Central PMCID: PMCPMC2132754.

55. Ben Q, An W, Fei J, Xu M, Li G, Li Z, et al. Downregulation of L1CAM inhibits proliferation, invasion and arrests cell cycle progression in pancreatic cancer cells in vitro. Exp Ther Med. 2014;7(4):785–90. Epub 2014/03/25. doi: 10.3892/etm.2014.1519. PubMed PMID: 24660028; PubMed Central PMCID: PMCPMC3961134.

56. Schafer H, Struck B, Feldmann EM, Bergmann F, Grage-Griebenow E, Geismann C, et al. TGF-beta1-dependent L1CAM expression has an essential role in macrophage-induced apoptosis resistance and cell migration of human intestinal epithelial cells. Oncogene. 2013;32(2):180–9. Epub 2012/02/22. doi: 10.1038/onc.2012.44. PubMed PMID: 22349829.

57. Basyoni MM, Rizk EM. Nematodes ultrastructure: complex systems and processes. J Parasit Dis. 2016;40(4):1130–40. Epub 2016/11/24. doi: 10.1007/s12639-015-0707-8. PubMed PMID: 27876901; PubMed Central PMCID: PMCPMC5118333.

58. Lee DL. The biology of nematodes. London: Taylor & Francis; 2002. xii, 635 p. p.

59. Attout T, Babayan S, Hoerauf A, Taylor DW, Kozek WJ, Martin C, et al. Blood-feeding in the young adult filarial worms Litomosoides sigmodontis. Parasitology. 2005;130(Pt 4):421–8. PubMed PMID: 15830816.

60. McGonigle S, Yoho ER, James ER. Immunisation of mice with fractions derived from the intestines of Dirofilaria immitis. Int J Parasitol. 2001;31(13):1459–66. PubMed PMID: 11595233.

61. Avery L, You YJ. C. elegans feeding: The C. elegans Research Community; 2012. WormBook:[Available from: http://www.wormbook.org.

62. Munn EA, Munn PD. Feeding and Digestion. In: Lee DL, editor. The Biology of Nematodes. London: Taylor & Francis; 2002. p. p. 211–33.

63. Scott AL, Ghedin E. The genome of Brugia malayi - all worms are not created equal. Parasitol Int. 2009;58(1):6–11. Epub 2008/10/28. doi: S1383-5769(08)00097-4 [pii] 10.1016/j.parint.2008.09.003. PubMed PMID: 18952001; PubMed Central PMCID: PMC2668601.

64. Newton SE, Morrish LE, Martin PJ, Montague PE, Rolph TP. Protection against multiply drug-resistant and geographically distant strains of Haemonchus contortus by vaccination with H11, a gut membrane-derived protective antigen. Int J Parasitol. 1995;25(4):511–21. Epub 1995/04/01. PubMed PMID: 7635627.

65. Munn EA. Rational design of nematode vaccines: hidden antigens. Int J Parasitol. 1997;27(4):359–66. Epub 1997/04/01. PubMed PMID: 9184927.

